# A gut sensor for sugar preference

**DOI:** 10.1101/2020.03.06.981365

**Authors:** Kelly L. Buchanan, Laura E. Rupprecht, Atharva Sahasrabudhe, M. Maya Kaelberer, Marguerita Klein, Jorge Villalobos, Winston W. Liu, Annabelle Yang, Justin Gelman, Seongjun Park, Polina Anikeeva, Diego V. Bohórquez

## Abstract

Animals innately prefer caloric sugars over non-caloric sweeteners. Such preference depends on the sugar entering the intestine.^1–4^ Although the brain is aware of the stimulus within seconds,^5–8^ how the gut discerns the caloric sugar to guide choice is unknown. Recently, we discovered an intestinal transducer, known as the neuropod cell.^9,10^ This cell synapses with the vagus to inform the brain about glucose in the gut in milliseconds.^10^ Here, we demonstrate that neuropod cells distinguish a caloric sugar from a non-caloric sweetener using the electrogenic sodium glucose co-transporter 1 (SGLT1) or sweet taste receptors. Activation of neuropod cells by non-caloric sucralose leads to ATP release, whereas the entry of caloric sucrose via SGLT1 stimulates glutamate release. To interrogate the contribution of the neuropod cell to sugar preference, we developed a method to record animal preferences in real time while using optogenetics to silence or excite neuropod cells. We discovered that silencing these cells, or blocking their glutamatergic signaling, renders the animals unable to recognize the caloric sugar. And, exciting neuropod cells leads the animal to consume the non-caloric sweetener as if it were caloric. By transducing the precise identity of the stimuli entering the gut, neuropod cells guide an animal’s internal preference toward the caloric sugar.

## Main Text

The cephalic senses guide our decision to eat. But what happens next, inside the gut, is essential for our eating preferences.

As early as 1952, it was known that animals prefer the side of a T-maze if rewarded by nutrients delivered directly into the stomach.^1^ Although the responses depend on the nutrient’s caloric value,^7,8,11^ how this value is signaled by the gut epithelium to drive preference is unknown. Of all macronutrients, sugars and available analogs have the most defined sensory properties. For instance, sucrose –a compound sugar made of D-glucose and fructose– contains both taste and calorie, whereas non-caloric sweeteners, like sucralose, carry only taste. Animals have an innate preference for sucrose over non-caloric sweeteners, even in the absence of taste.^3,12–14^ Such preference depends on the sugar entering the intestine.^2,3,15,16^ But slow acting intestinal hormones cannot account for the effect,^17^ because the brain perceives stimuli from nutrients entering the small intestine within seconds.^6,8,10^ Thus, a fast intestinal sensor must exist to steer the animal towards the caloric sugar.

### A gut sense for sweets

Intestinal signals arising from the lumen are relayed to the brain via the vagus nerve.^6,10,11^ The vagus responds within seconds to an intraluminal stimulus of sucrose,^6,10,18^ but the response to other sugars, including non-caloric sweeteners, is unclear. We therefore tested vagal firing responses to a panel of sugars: sucrose, D-glucose, fructose, galactose, maltodextrin, alpha-methylglucopyranoside (α-mgp), saccharin, acesulfame-K, and sucralose. All stimuli were perfused at physiological concentrations^10^ through the proximal duodenum, bypassing gustatory or gastric activation, while nerve responses were recorded using two electrodes placed at the cervical vagus (Fig. 1a). Almost all sugars elicited a significant increase in vagal firing rate within seconds (p<0.001, n≥5; Fig. 1b, Extended Data Fig. 1a-b). But fructose did not (Extended Data Fig. 1c-d). Unlike D-glucose, it diffuses passively through the epithelium and does not have a clear reinforcing effect.^19^ All responses were confined to the small intestine, the primary site for sugar absorption^20^, which correlates with post-ingestive rewarding effects^4^ (Extended Data Fig. 2a-c).

**Fig. 1.**
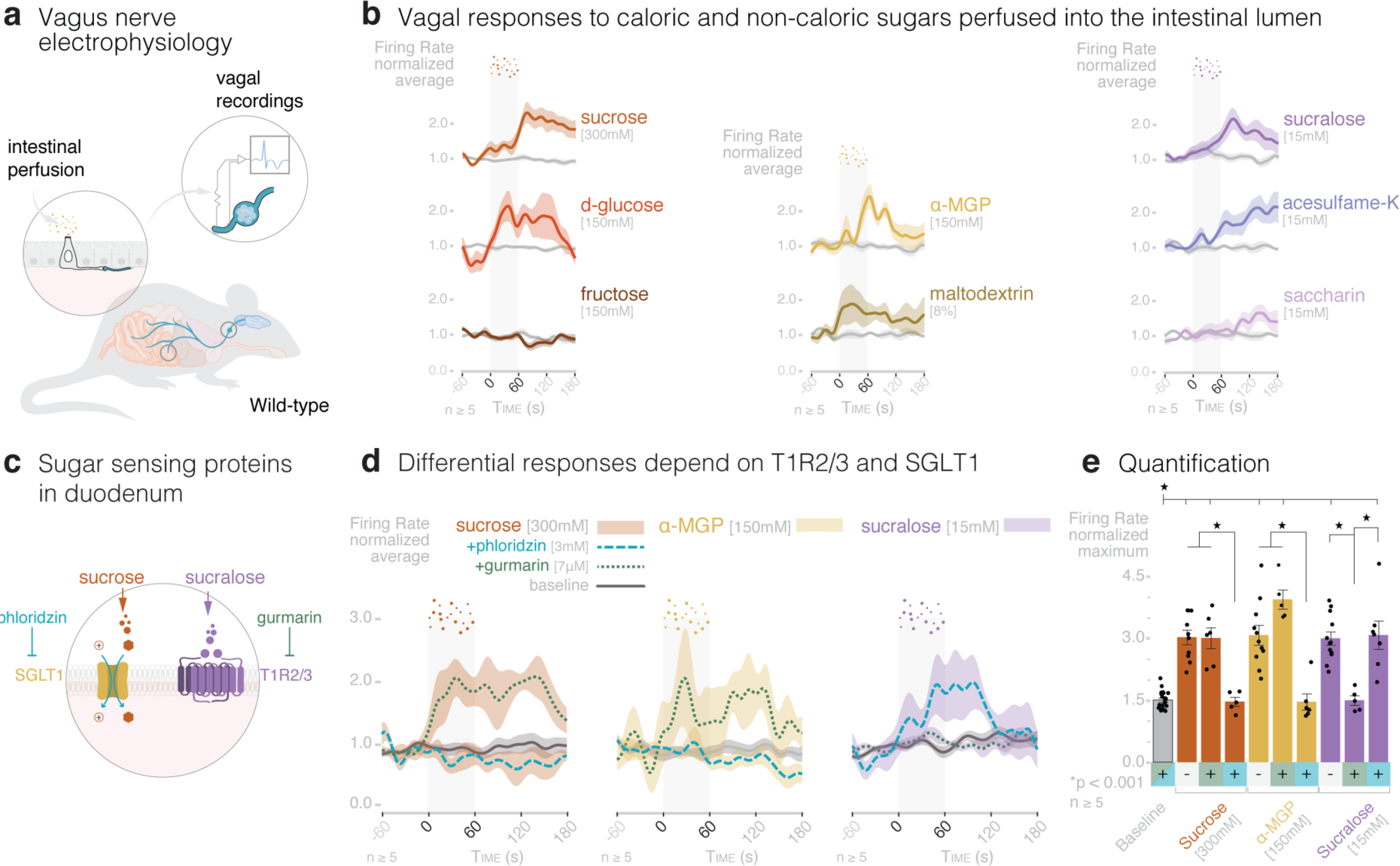
Intraluminal sugars are represented in the vagus via the activation of sweet taste receptors (T1R2/3) or sodium-glucose co-transporter 1 (SGLT1). **a**, Model: sugars are perfused into the duodenal lumen while nerve activity is recorded from the cervical vagus of anesthetized wild-type mice. **b**, The vagus responds significantly (p<0.05) to sucrose [300mM], D-glucose [150mM], maltodextrin [8%], α-mgp [150mM, sucralose [15mM], and acesulfame K [15mM] (n≥5 wild-type mice per group; quantified in Extended Data 1). Fructose [150mM] and saccharin [15mM] responses are not significant (p>0.05). **c**, Model: duodenal epithelial cells are known to express SGLT1 and T1R2/3 which are inhibited by phloridzin and gurmarin, respectively. **d**, Vagal responses to intra-duodenal sucrose [300mM] and α-MGP [150mM] are inhibited by SGLT1 inhibitor phloridzin [3mM] and to sucralose [15mM] by the T1R2/3 inhibitor gurmarin [7μM]. **e**, Quantification of e; (n≥5 mice per group; *p<0.0001, ANOVA with post-hoc Tukey’s HSD test). Gray vertical bars indicate infusion period. Shaded regions on traces or error bars indicate SEM.

Intestinal epithelial cells absorb glucose through active transport mediated by the sodium glucose co-transporter SGLT1. This electrogenic channel uses a sodium gradient to transport D-glucose into the cell. Thus, SGLT1 acts as a gate for D-glucose.^21,22^ Intestinal epithelial cells also express sweet taste receptors, specifically T1R2 and T1R3.^23^ We found that the vagal response to sucralose [15mM] was abolished when these taste receptors were inhibited with gurmarin [7μM]^24^ (Fig. 1c-e), but the response to sucrose [300mM] or α-mgp [150mM] remained intact (Fig. 1c-e). α-mgp is a synthetic sugar that once inside the cell is not metabolized but, like D-glucose, α-mgp enters the cell through SGLT1 (Extended Data Fig. 2d).^21^ As such, inhibiting SGLT1 with phloridzin [3mM]^25^ abolished the response to both α-mgp and sucrose (Fig. 1c-e). Phloridzin did not affect the vagal response to the luminal sucralose stimulus (Fig. 1c-e). The sweetness and caloric value of sugars are therefore represented in the vagus by two distinct signals emanating from the intestinal epithelium.

### The sensor

Within the epithelium of the intestine, there is sensory transducer. This is the neuropod cell – an electrically excitable cell that synapses with nerves.^5,10,26,27^ Previously, we discovered that intestinal neuropod cells transduce a D-glucose stimulus to vagal neurons in milliseconds.^10^ But it is unknown if this neuroepithelial circuit can distinguish the sweetness and caloric value of sugars.

Vagal nodose neurons alone do not respond to sugars. 58 of 59 dissociated neurons showed no calcium transients in response to D-glucose [20mM], maltodextrin [1%], or sucralose [2mM]. All neurons responded to KCl [50mM], confirming their viability (n=59 neurons; Extended Data Fig. 3a). A supplementary single cell sequencing analysis showed that nodose neurons lack the expression of *Tas1r2, Tas1r3*, and the SGLT1 transcript *Slc5a1* (N≥5 left and right nodose ganglia, Extended Data Fig. 3b). T1R2, T1R3 and SGLT1 do, however, colocalize with intestinal epithelial cells immunoreactive for the cholecystokinin neuropeptide.^23,28^

Intestinal neuropod cells are labeled by the promoter for cholecystokinin.^5,10^ This neuropeptide only participates in the prolonged vagal responses to D-glucose minutes after the initial stimulus.^10^ To determine if neuropod cells rapidly sort sugar stimuli, we used optogenetics. Using Cre-loxP recombination, we bred a CckCRE_Halo mouse in which neuropod cells expressed the chloride pump halorhodopsin (Fig. 2a). When triggered by 532nm light, halorhodopsin hyperpolarizes the cell membrane, silencing electrically excitable cells instantly. In CckCRE_Halo mice, vagal responses to luminal sucrose, α-mgp, and sucralose were abolished in the presence of 532nm light (n ≥ 5, p<0.001; Fig. 2b-c). Vagal responses remained intact in the presence of the control 473nm light.^10^ Thus, neuropod cells are necessary to transduce both caloric and non-caloric sugar stimuli.

**Fig. 2.**
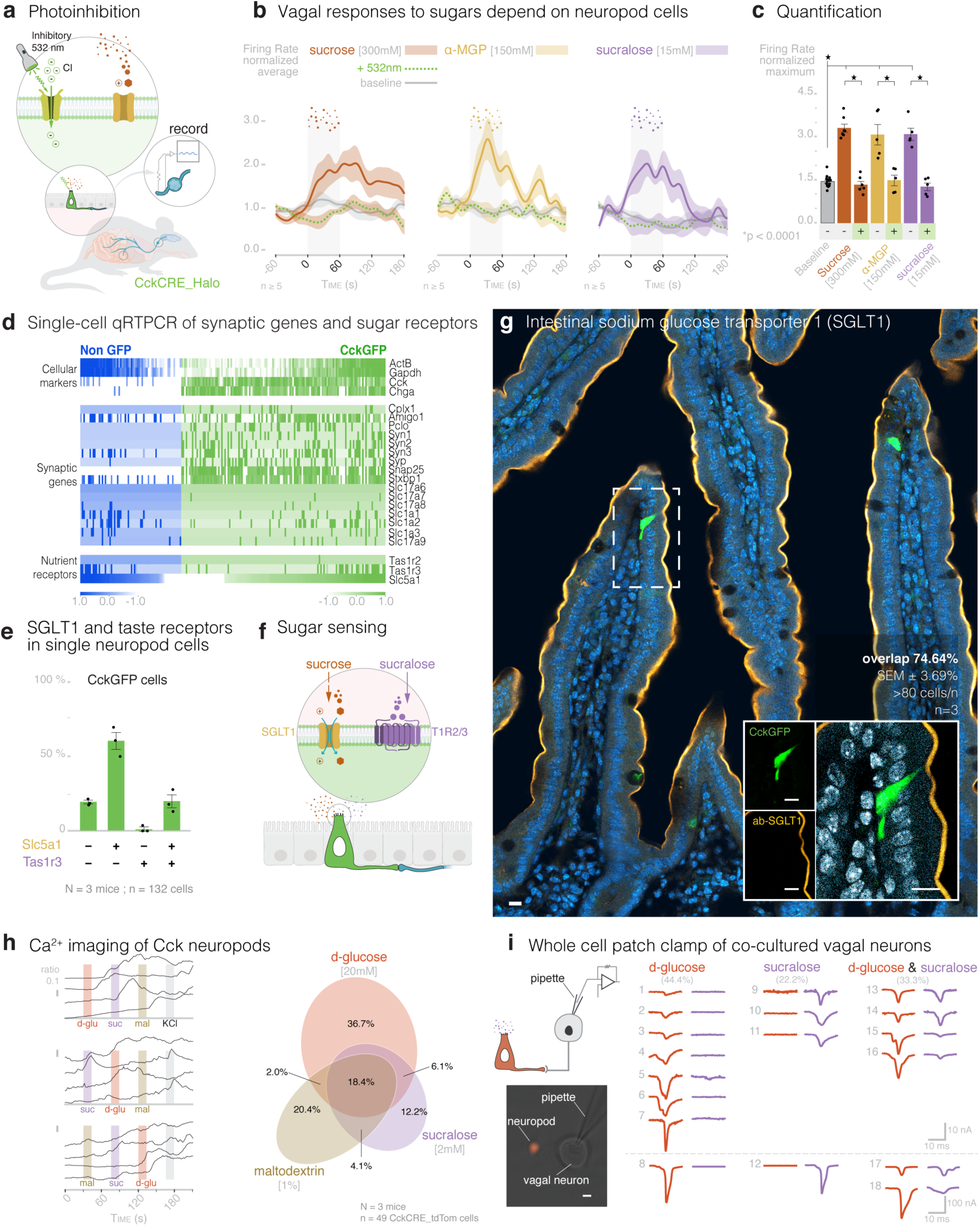
Neuropod cells sort caloric sugars from non-caloric sweeteners. **a**, Model: intraluminal 532nm laser activates Halorhodopsin chloride channels to silence duodenal neuropod cells in CckCRE_Halo mice. **b**, Silencing duodenal CckCRE_Halo cells eliminates vagal responses to sucrose [300mM], α-mgp [150mM], and sucralose [15mM). **c**, Quantification of b (n≥5 mice per group; *p < 0.0001, ANOVA with post-hoc Tukey’s HSD test). Gray vertical bars indicate infusion period. **d**, Single cell qRT-PCR of CckGFP cells and non-GFP intestinal epithelial cells. Compared to non-GFP epithelial cells (n=66), CckGFP cells (n=132) overexpress pre-synaptic genes *Cplx1, Amigo1, Pclo, Syn1, Syn3, Syp, Snap25, Stxbp1* (N=3 mice; q=0.01). **e**, Of 132 CckGFP cells, 19.1 ± 1.2% express transcripts for neither *Slc5a1* (SGLT1) nor *Tas1r3* (T1R3), 60.1 ± 5.7% for only *Slc5a1*, 1.2 ± 1.2% for only *Tas1r3*, and 19.6 ± 4.3% for both, (n=3 mice). **f**, Model: neuropod cells express SGLT1 and T1R2/3 to sense sugars. **g**, ∼3 out of 4 CckGFP cells (green) in the duodenum express SGLT1 (yellow) (n=80-100 cells per mouse, N=3 mice). Scale bars, 10μm. **h**, Intestinal CckCRE_tdTomato cells loaded with Fluo-4/FuraRed respond to D-glucose [20mM], sucralose [2mM], and maltodextrin [1%]. *Left-*representative traces; *right-*Venn diagram. (n=3 mice, n=49 cells). **i**, *Left-*Patch-clamp electrophysiology of neurons in co-culture with CckCRE_tdTomato cells (*top-*model; *bottom-*image, bar=10μm). *Right-*Of 18 co-culture pairs, excitatory post-synaptic potentials were recorded in neurons in response to D-glucose (20mM, 44.4%), to sucralose (2mM, 22.2%), and to both (33.3%). Shaded regions on traces and error bars indicate SEM.

### Taste and calorie sorted

Receptors for both caloric and non-caloric sugars were expressed in individual CckGFP cells. *Tas1r2* was negligible, *Tas1r3* alone was in 1.2% (±1.2), *Tas1r3* and the SGLT1 transcript *Slc5a1* were in 19.6% (±4.3), and the SGLT1 transcript *Slc5a1* alone was in 60.1% (±5.7) (N = 3 mice, n = 132 CckGFP cells; Fig. 2d-f). Accordingly, SGLT1 was immunoreactive in 74.6% (±3.7) of CckGFP cells (N = 3 mice; n ≥ 80 cells per mouse; Fig. 2g). Moreover, compared to other intestinal epithelial cells, *Slc5a1* positive CckGFP cells had significantly increased expression of the pre-synaptic genes *Efnb2* (fold change 81.6) and *Cask* (fold change 30.2), and the synaptic adhesion genes *Pvrl1* (fold change 31.05) and *Pvrl2* (fold change 35.2) (n = 104 *Slc5a1*+ CckGFP+ cells, n = 28 *Slc5a1-*CckGFP+ cells; *p<0.0001) (Fig. 2d). Synaptic molecules are a feature of fast acting neuropod cells.^5,10^

We determined if individual neuropod cells respond to both sweet and caloric sugars. Using calcium imaging, we recorded responses from individual Cck_tdTomato cells. Stimuli were applied in sequence and alternated in replicate experiments to account for potential priming or inhibitory effects. Cell viability was confirmed using KCl [50mM]. Responses were as follows: 36.7% responded to D-glucose alone [20mM], 20.4% to maltodextrin alone [1%], 12.2% to sucralose alone [2mM], and 18.4% to all three sugars (N = 3 mice, n = 49 cells; Fig. 2h). To determine if these responses are represented in connected vagal neurons in the absence of other cell types, we co-cultured Cck_tdTomato cells with nodose neurons. In vitro, neuropod cells synapse with vagal neurons.^5,10^

We recorded activity from connected neurons using voltage-clamp electrophysiology. D-glucose and sucralose stimuli were applied to the bath solution in alternating order (Fig. 2i). We observed excitatory post-synaptic currents in connected neurons as follows: 44.4% to D-glucose alone, 22.2% to sucralose alone, and 33.3% to both sucrose and sucralose (n = 18 pairs, Fig. 2i). Peak current in the connected neurons was not statistically different for either stimulus (Extended Data Fig. 3d), suggesting that two separate paths, of similar synaptic strength, are elicited by the sweet and caloric stimuli sensed by neuropod cells.

### The messengers

D-glucose causes the release of glutamate from neuropod cells.^10^ These cells express transcripts for the vesicular glutamate transporters *Slc17a7, Slc17a8* and the synaptic glutamate transporters *Slc1a1, Slc1a2*, and *Slc1a3* (Fig. 2d). In addition, vagal neurons express transcripts for both metabotropic and ionotropic glutamate receptors (Extended Data Fig. 3e). But it remained to be determined whether both sweet taste and caloric sugars trigger glutamate release (Fig. 3a).

**Fig. 3.**
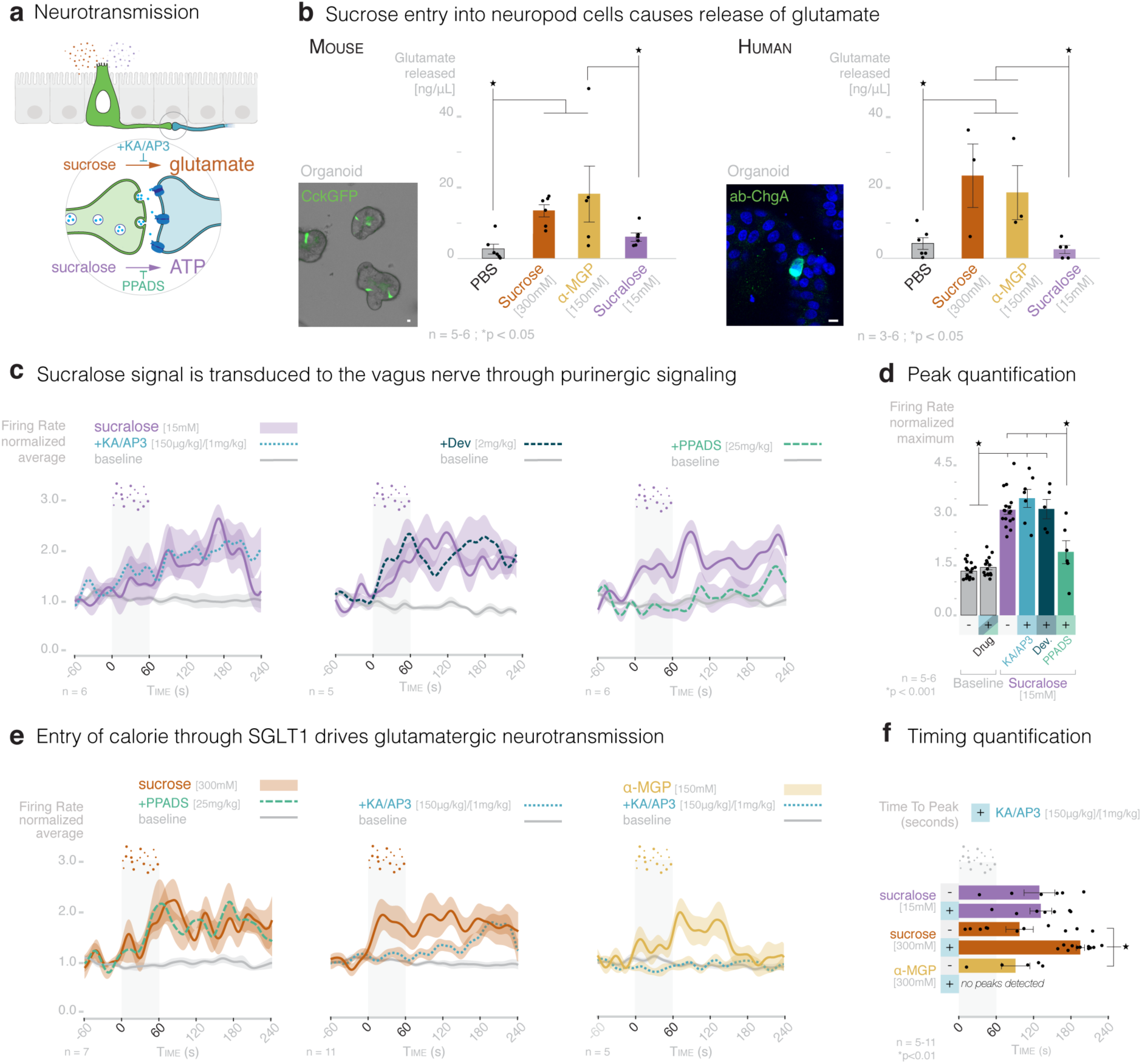
Sucralose leads to ATP release, whereas the entry of sucrose via SGLT1 stimulates glutamate release. **a**, Model: neuropod cells transduce caloric and non-caloric sugar signals to the vagus nerve using differential neurotransmitter release. **b**, *Left*-In CckGFP mouse intestinal organoids (CckGFP, green), sucrose [300mM] and αMGP [150mM] elicit significant glutamate release compared to PBS control, while sucralose [15mM] does not (n=5-6 plates, N=3 mice, *p<0.05 by Student’s t-test). *Right*-Human duodenal organoids contain Chromogranin-A+ cells (ChgA, green) and release glutamate in response to sucrose and αMGP, but not to sucralose or PBS control (n=3-6 plates, N=1 human sample, *p<0.05 by Student’s t-test). **c**, The rapid vagal response to sucralose [15mM] is not attenuated by inhibition of ionotropic/metabotropic glutamate receptors with KA/AP3 [150μg/kg]/[1mg/kg] or by inhibition of Cck-A receptors with devazepide [2mg/kg]; rather, it is ATP-dependent (attenuated by non-selective P2 purinoceptor inhibition with PPADS [25mg/kg]). **d**, Quantification of c (n=5-6 mice, *p<0.001 ANOVA with post hoc Tukey’s HSD test). **e**, While inhibition of P2 purinoreceptors does not affect the response to sucrose, the fast vagal response to sucrose [300mM] and the entire response to α-MGP [150mM] are abolished by glutamate receptor inhibition with KA/AP3. **f**, Glutamate receptor inhibition prolonged the time to peak vagal firing rate in response to sucrose but not sucralose. No increase in vagal firing rate was detected for α-MGP after glutamate receptor inhibition (n=5-11 mice, *p<0.01 ANOVA with post hoc Tukey’s HSD test). Shaded regions on traces or error bars indicate SEM. Scale bars, 10μm.

We tested glutamate release by culturing organoids from mouse proximal small intestine^29^ and human duodenum.^30^ In both mouse and human organoids, sucrose [300mM] and α-mgp [150mM] elicited a significant release of glutamate compared to PBS (N = 3 mice, n ≥ 5 organoid wells in triplicate per stimuli, p<0.05, Fig. 3b-left; N = 1 human patient donor, n ≥ 3 organoid wells in triplicate per stimuli, p<0.05, Fig. 3b-right). Sucralose [15mM] did not stimulate glutamate release (Fig. 3b; Extended Data Fig. 4a).

In fact, the vagal response to sucralose remained intact when glutamate receptors were blocked using an intraluminal injection of AP3 [1mg/kg] plus kynurenic acid [150μg/kg] (Fig. 3c-d). Although vagal neurons express the cholecystokinin A receptor, the vagal response to sucralose remained intact even when blocking cholecystokinin signaling with devazepide^17^ [2mg/kg] (Fig. 3c-d; Extended Data Fig. 3e, 4b). In other sensory transducers, like taste receptor cells in the tongue, sweet stimuli are transduced onto to afferent nerves using vesicular ATP.^31^ We observed that vagal nodose neurons express ATP purinergic receptors and CckGFP cells express the *Slc17a9* transcript for the vesicular nucleotide transporter (Extended Data Fig. 3e; Fig. 2d). Indeed, by inhibiting purinergic receptors with PPADS [25mg/kg], vagal responses to sucralose were significantly attenuated. This indicates that sweet sucralose is transduced by neuropod cells using ATP as a neurotransmitter (p<0.05, Fig. 3c-d; Extended Data Fig. 4e-f).

### It is the entry of the sugar

Inhibiting ATP neurotransmission did not affect the vagal response to sucrose or α-mgp (Fig. 3e-f, Extended Data Fig.4d-f). Instead, the entire α-mgp response and the first 120 seconds of the sucrose response were abolished by blocking glutamate neurotransmission (Fig. 3e-f; Extended Data Fig. 4c). If the α-mgp response completely depends on glutamatergic neurotransmission and α-mgp enters the cell through SGLT1, then the entry of the sugar through SGLT1 is sufficient to drive glutamatergic neurotransmission between neuropod cells and vagal neurons. We wondered if this neuropod cell mechanism is responsible for the animals’ innate preference of caloric sugars over non-caloric sweeteners.

### Timing sugar preferences

To determine the timing of the behavioral response in seconds, without perturbing the animal’s natural behavior, we optimized an automated phenotyping system called PhenoMaster (Fig. 4a-b). Each PhenoMaster cage holds one mouse and two gated bottles that are constantly weighed with a time resolution of 5 seconds. We gave the mouse a choice between iso-sweet sucrose [300mM] and sucralose [15mM]. When the gates opened, experienced mice showed a robust and significant preference for sucrose within ∼125 seconds (p < 0.05, n = 9 mice; Fig. 4b). Now we needed a method to record the mouse’s preferences while its neuropod cells are rapidly “switched on or off”.

**Fig. 4.**
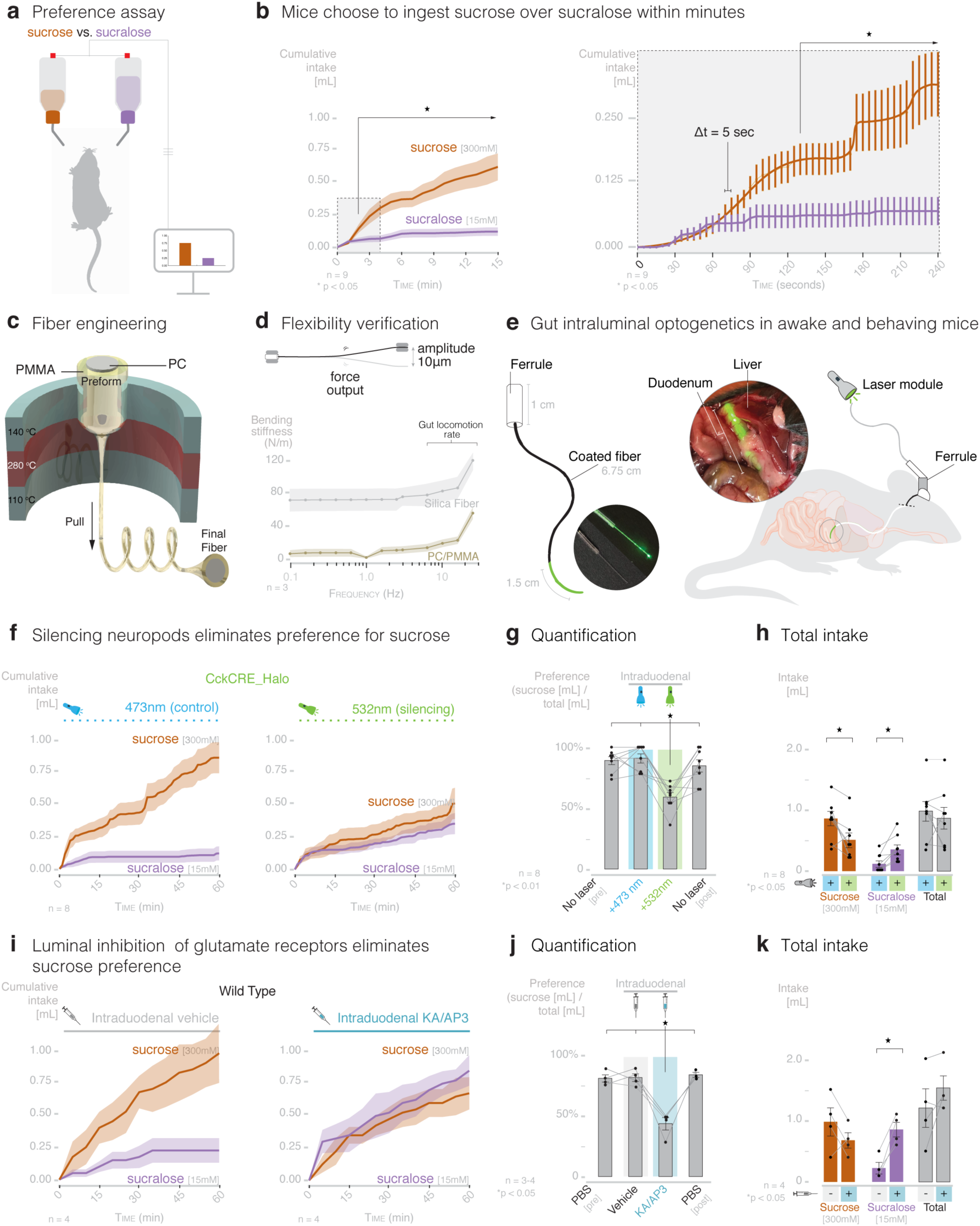
Intestinal neuropod cells drive choice for sucrose over sucralose. **a**, Model: mice choose between sucrose [300mM] or sucralose [15mM] while consumption is recorded every 5 seconds. **b**, Within 2.08 minutes, wild-type mice develop sucrose preference (n=9 mice, *p<0.05, repeated-measures ANOVA with post-hoc paired t-test). **c**, Schematic: thermal drawing process for the flexible PC/PMMA waveguide fiber. Application of controlled stress and heat reduces diameter ∼40 fold. **d**, Flexibility of standard silica and PC/PMMA fiber was measured by a dynamic mechanical analyzer in single cantilever mode with a displacement amplitude of 10 μm at physiologic frequencies (n = 3 fibers, shaded regions represent SD)**. e**, Four weeks after intraduodenal implantation, the flexible fiberoptic continues to illuminate. **f**, Compared to 473nm control laser, silencing duodenal neuropod cells with 532nm laser in CckCRE_NpH3 mice eliminates sucrose preference. **g**, Preference quantified at one hour with no (pre/post), 473nm (control), and 532nm (silencing) laser. **h**, Silencing neuropod cells decreases sucrose and increases sucralose consumption but does not affect total consumption. **i**, In wild-type mice, intraduodenal perfusion of KA/AP3 [5ng/0.1μg in 0.4mL over 1 hour] to inhibit local glutamate receptors eliminates sucrose preference. **j**, Preference quantified at 1 hour with PBS (pre/post), vehicle (PBS+NaOH, pH 7.4), and glutamate receptor blockers KA/AP3. **k**, Inhibiting glutamate receptors increases sucralose consumption without significantly affecting sucrose or total consumption (for f-h: n=8; for i-k: n=4; *p<0.05, repeated-measures ANOVA with post-hoc paired t-test). Shaded regions or error bars indicate SEM unless otherwise indicated.

In the brain, the contribution of specific neural circuits to behaviors has been widely explored using light-sensitive opsins triggered with laser light delivered via fiberoptic cables. But in the gut, the rigid silica fiberoptics suitable for the mouse brain puncture and perforate the delicate intestinal lumen. We learned rather quickly that flexibility is a must.

### The illumenator

To illuminate the intestines of awake and behaving mice, we envisioned a flexible fiberoptic waveguide fiber with the following properties: (1) thin in diameter for minimal footprint within the intestinal lumen, 2) low optical loss coefficient, and 3) durable for months when flexed and bent inside the churning gut. We engineered a template of a poly-methyl methacrylate (PMMA) cladding layer around an optical core of polycarbonate (PC). The template was consolidated using thermal drawing process^32,33^ and pulled at 270°C into a final flexible fiberoptic of only 230μm in diameter (Fig. 4c; Extended Data Fig. 5a). Compared to rigid silica fibers, the flexible fiberoptic bends and does not pierce through even a soft layer of 1.5% agarose (Extended Data Fig. 5b, Video 1). The device efficiently transmits light and tolerates rapid bending at 10Hz, which is above the physiological frequency of gut motility (Fig. 4d; Extended Data Fig. 5c-e). We called the device the illumenator.

Prior to implantation, we enclosed the illumenator in an opacified microrenathane sheath to restrict the delivery of the light only to a portion of the tip end (Fig. 4e). When implanted, this feature allowed us to target the light only to the first 1.5cm of the mouse small intestine. We confirmed the device’s durability by implanting it in the intestine of CckCRE_Halo mice. Four weeks later, a 532nm light stimulus emitted from the device was sufficient to abolish normal vagal responses to sucrose perfused into the duodenum (Extended Data Fig.5f).

We implanted mice with the illumenator to test sugar preferences (Fig. 4e). Each mouse was acclimated for neophobia, tested for side preference, and the location and power of the implanted device were corroborated at the end of the study (see methods). On each experimental day, implanted mice were given the choice between sucrose and sucralose for 1 hour while receiving laser stimulation (1min on/2min off, 5V, 40Hz, 20% duty cycle). Sucrose and sucralose intake were recorded at 5 second intervals. Locomotor activity was recorded throughout the study, and chow and water intake were recorded for 24 hours following the assay to control for malaise.

### Neuropods drive sugar preference

In CckCRE_Halo mice, silencing duodenal neuropod cells with 532nm light completely abolished the innate preference for sucrose. In the presence of 532nm light, negative littermates showed 90.8% (± 3.74) sucrose preference, whereas CckCRE_Halo mice only showed 58.9% (± 3.92) (n ≥ 5, p < 0.01; Fig. 4f-g, Extended Data Fig. 6d-e). Silencing neuropod cells did not affect chow or water intake in the following 24 hours, or locomotor activity during the assay, indicating no malaise effects (Extended Data Fig. 6a-c). Nor did it affect the total consumption of liquid during the hour test. Instead, it reduced sucrose intake and increased sucralose intake (p < 0.05; Fig. 4h, Extended Data Fig. 6f). Sucrose preference was not affected by blocking cholecystokinin receptors with intraperitoneal devazepide (Extended Data Fig. 7a-d).^17^

We wondered if glutamate is the transmitter for sucrose preference. A catheter was implanted into the mouse’s duodenal lumen to deliver a low dose of the glutamate receptor blockers, AP3 plus kynurenic acid [0.1µg/5ng in 0.4mL]. Mice underwent the same inclusion analysis as those implanted with the illumenator. Sucrose preferences in mice receiving a vehicle control was 82.4% (± 3.20), whereas in mice receiving intraduodenal glutamatergic inhibitors, sucrose preference was significantly reduced to 44.0% (± 5.19) (n = 4, p < 0.05; Fig. 4i-j). No malaise effects were observed. Blocking glutamatergic neurotransmission also increased sucralose intake without significantly changing total intake (Fig. 4k). In the absence of the neuropod cell glutamatergic input, the animal could not distinguish the caloric sugar from the non-caloric sweetener.

If so, could exciting neuropod cells increase sucralose intake? We bred mice in which neuropod cells express channelrhodopsin 2 (ChR2). ChR2 depolarizes electrically excitable cells instantly if stimulated with 473nm light. In this test, intake was paired to laser stimulation (0.01mL intake caused 5 seconds stimulation; 5V, 40Hz, 20% duty cycle). When presented with one bottle of sucralose, CckCRE_ChR2 mice significantly increased their intake if their duodenal neuropod cells were excited with 473nm light. Exciting neuropod cells increased the mouse’s intake of sucralose to its innate level of sucrose (n = 4, p < 0.05; Extended Data Fig. 8a-c). Thus, neuropod cells drive sugar intake and preference.

### A gut choice

In his classic book *Behave*, the neuroendocrinologist Robert Sapolsky states “What occurred in the prior seconds to minutes that triggered the nervous system to produce the behavior, this is the world of sensory stimuli, much of it sensed unconsciously.”^34^ By sensing the entrance of sugar, neuropod cells have the responsibility to advise the nervous system to steer the animal towards the sugar with the caloric value, even before consciousness has settled. This work is a foundation to understand how the nutritive value of other ingested macronutrients, and perhaps micronutrients, are sensed by the gut to aid the brain in choosing what to put in our mouth.

## Acknowledgements

The authors wish to thank Mr. Peter Weng, Ms. Marcia Montoya Gomez, Mr. Bradley Barth, Mr. Brayan Florentino, and Ms. Evie Freel for their contributions. We thank Mr. Carlton Anderson and Ms. Gabrielle Kelly in the CGIBD Advanced Analytics Core for assistance in single cell qPCR experiments. Our sincere appreciation is expressed to the staff of the Duke Light Microscopy Core, Flow Cytometry Core, and Department of Animal Research. Funding: HHMI Medical Research Fellowship to K.L.B., T32 DK007568 to M.M.K., Hartwell Postdoctoral Fellowship to L.R., F30 DK122712 to W.W.L, and DP2 MH122402, R21 AT010818, and Duke NUS Pilot Research Grant to D.V.B.

## Author Contributions

K.L.B. optimized and performed flexible fiber implantation surgery and behavior experiments and performed all vagal cuff recordings, single cell qPCR experiments, optogenetic behavioral studies, and associated data analysis. L.E.R. optimized and performed pharmacologic and channelrhodopsin2 behavior experiments. A.S., S.K., and P.A. designed and fabricated the flexible fiberoptic. M.M.K. performed all single cell calcium imaging, co-culture patch clamp electrophysiology experiments, and associated data analysis. M.E.K. cultured and maintained all organoids. M.E.K., K.L.B., and A.Y. optimized and performed glutamate release assays. K.L.B. and W.W.L. optimized the behavioral phenotyping system. W.W.L. analyzed single cell RNA sequencing data. J.V. planned and performed all animal breeding, mouse colony management, genotyping, and quality control. J.G. performed immunohistochemistry and quantification. K.L.B. and D.V.B. planned experiments and composed figures. D.V.B. conceptualized project, supervised the research, and wrote the final manuscript.

### Competing interests

Some of the findings in this manuscript have been used to file a provisional patent application. D.V.B. is a member of the scientific advisory board of Holobiome Inc. on a fee-for-service basis. No other competing interests are declared.

### Data and materials availability

All data is available in the manuscript, the supplementary materials, or upon request from the authors.

## Extended Data

## Methods

### Mouse strains

All experiments on mice were performed following approval by the Institutional Animal Care and Use Committee at Duke University Medical Center under the protocol A280-18-12. Mice were group housed in Duke University’s Division of Laboratory Animal Resources, where they were kept on a 12-hour light-dark cycle (0700-1900) with access to water and standard mouse chow (Purina 5001) *ad-libitum*, unless otherwise indicated. Male and female adult mice aged 4-20 weeks were used in all experiments. The mouse strains, source, background, and stock number used to breed experimental mice are listed in the table below. The following experimental mouse strains were purchased/received/bred in-house and used directly: C57Bl6/J, Swiss Webster, and CckGFP. The following double transgenic mouse strains were bred in-house: CckCRE_tdTomato, CckCRE_Halo-YFP, and CckCRE_ChR2-tdTomato.

| Mouse genotype | Source | Background | Stock Number |
| --- | --- | --- | --- |
| Wild type | Jackson Lab | C57BL/6J | 000664 |
| Wild type | Charles River | Swiss Webster | 024 |
| CckGFP | Courtesy of Liddle Laboratory | Swiss Webster | Wang *et al.*, (2010)<sup>35</sup> |
| CckCRE | Jackson Lab | C57BL/6J | 012706 |
| LSL_tdTomato | Jackson Lab | C57BL/6J | 007914 |
| LSL_Halo-YFP | Jackson Lab | C57BL/6J | 014539 |
| LSL_ChR2-tdTomato | Jackson Lab | C57BL/6J | 012567 |

\*LSL = loxP-STOP-loxP cassette

### Human samples

Human duodenal samples were obtained from the Duke University Medical Center Biorepository and Precision Pathology Center (BRPC) under the Institutional Review Board (IRB) protocol Pro00035974 via anonymous tissue release. All samples were deidentified and all links to additional patient information were broken prior to receipt of fresh surgical specimens. Following surgical extraction, samples were placed in sterile PBS and stored at 4°C prior to crypt dissociation.

### Organoid culture

#### Murine organoids

Mouse organoids were cultured from CckGFP mice (n = 3) per Sato *et al.* (2009).^36^ Briefly, the proximal one third of the small intestine was flushed with cold PBS, opened lengthwise and cut into ∼1cm pieces. Tissue pieces were incubated with 1.5mM EDTA on ice for 25 minutes then 37°C for 15 minutes. Crypts were detached by shaking in cold PBS, pelleted at 100xg and resuspended in Matrigel (Corning #356231). Crypts were plated in 50µl mounds in 24 well plates and maintained in organoid culture media containing 1x Glutamax (Gibco), 10mM HEPES (Gibco), 200U/ml Penicillin-Streptomycin (Gibco), 1x N2 supplement (Gibco), 1x B27 supplement (Gibco), 1mM N-acetylcysteine (Sigma), 50ng/ml EGF (Peprotech), 100ng/ml Noggin (Peprotech), and 10% R-spondin conditioned media (produced in house using Trevigen 3710-001-K cells) in Advanced DMEM/f12 (Gibco). 10μM Y-27632 (Enzo) was added to the culture media for initial plating.

#### Human organoids

Human organoids were cultured per Fujii *et al.* (2015).^30^ Human tissue was washed in PBS and the epithelial layer was dissected from submucosa and connective tissue and minced. All tips and tubes used were coated with FBS. Tissue pieces were washed in PBS until washes were clear then incubated in 5mM EDTA in PBS for 1 hour, on ice. Crypts were detached by shaking in cold PBS, pelleted at 100xg and resuspended in Matrigel (Corning #356231). Crypts were plated in 50µl mounds in 24 well plates and maintained in Human Intesticult (Stem Cell #06010) with 10μM Y-27632 at initial plating.

### Glutamate release assay

Murine or human small intestinal organoids were passaged and plated into a 96 well plate in 25μL Matrigel mounds in organoid culture media as above. The human organoid media contained 500ng/ml human r-spondin (Peprotech) and was supplemented with 500nM A-83-01 (Tocris) and 10nM leu-gastrin (Sigma) for differentiation with 10μM Y-27632 at passage.^37^ When mature morphology was achieved 3-7 days after passage, media was removed and organoids were washed in PBS twice for 5 minutes at RT. Organoids were then stimulated with 60μL of 300mM sucrose, 150mM alpha-methylglucopyranoside (α-MGP), 15mM sucralose, or PBS for 10 minutes at 37°C. Supernatant was collected, centrifuged for 10 minutes at 13,000xg to remove insoluble material, and stored at -20°C for up to 2 weeks. Glutamate concentration in samples was assessed using the Glutamate Release Assay Kit (Sigma). 30μL of each sample was mixed with buffer, glutamate enzyme mix, and developer following manufacturer’s protocol. Each experimental condition was run in triplicate on every plate. A glutamate standard was run for every plate. Control wells of sample and developer without enzyme mix were run in duplicate and included for each sample. 450nm absorbance was measured on a plate reader (Tecan Infinite 200 Pro). Nine reads were taken per well and averaged. The absorbance from the control was subtracted from each experimental sample absorbance for the corrected value. Glutamate amount and concentration was calculated using the standard curve.

### Vagus nerve recordings

Whole nerve recordings were performed in wild-type mice (n = 5-11 per group), CckCRE_Halo-YFP mice (n = 5 per group), and CckCRE_Halo-YFP mice following fiberoptic implantation (n = 4-5 per group). Whole nerve electrophysiology recordings of the cervical vagus nerve were performed as previously reported.^10^ A 20-gauge gavage needle with two connected tubes for PBS perfusion and stimulant delivery was surgically inserted through the stomach wall into the duodenum (for small intestine experiments) or the proximal colon distal to the cecum (for colon experiments). A perfusion exit incision was made at the ligament of treitz for the small intestine or just proximal to the rectum for colon. PBS was constantly perfused through the isolated intestinal region at ∼400μL per minute as a baseline and volume pressure control. Stimulation conditions were applied after recording 2 minutes of baseline activity. During nutrient stimulation conditions, PBS perfusion was continuous and 200μL of stimulant was perfused over 1 minute using a syringe pump (Fusion 200, Chemyx). All caloric and non-caloric sugar ligands were purchased from Sigma and dissolved in PBS. The final concentration for each infused nutrient is as follows: 8% maltodextrin, 150mM alpha-methylglucopyranoside (α-MGP), 15mM sucralose, 15mM acesulfame K (ace-K), 15mM saccharin, 300mM sucrose, 150mM d-glucose, 150mM galactose, 15mM-1M fructose. All sugar ligands were purchased from Sigma.

### Data Acquisition

Extracellular voltage was recorded as previously described.^10^ The raw data was analyzed using Spike Tailor, a custom MATLAB software (Mathworks) script.^10^ Spikes were detected using a threshold detected based on RMS noise. The firing rate was calculated using a Gaussian kernel smoothing algorithm in 200-ms bins.^38^

### Optogenetic inhibition

A standard stiff silica fiber optic cable (FT020, ThorLabs) was threaded through or along the gavage needle into the duodenal lumen. Laser stimulation was applied simultaneously with nutrient infusion. The laser was pulsed for 1 minute at 40Hz, 5V peak, and 20% duty cycle (473nm, 80mW laser, RGBlase; 532nm, 80mW laser, RBGlase).

### SGLT1/T1R2/3 inhibition

For apical receptor inhibition, the SGLT1 competitive inhibitor phloridzin dihydrate (Sigma) or the sweet taste receptor inhibitor (T1R2/3) gurmarin (Peptides International) were dissolved into 1M sucrose, 45mM sucralose, or 450mM α-MGP. Following recording of a pre-inhibitor response to the selected ligand, sucrose, sucralose, or α-MGP were perfused with the apical receptor inhibitors, for a final phloridzin concentration of 3mM^10^ and a final gurmarin concentration of 7μM.^39,40^ Each mouse received one only inhibitor.

### Neurotransmitter inhibition

The following neurotransmitter/neuropeptide blockers were used: CCK-A receptor antagonist devazepide ([2mg/kg] in 5% DMSO PBS; Sigma)^10,41^; cocktail of the ionotropic glutamate receptor antagonist kynurenic acid (KA) ([150μg/kg] in PBS, stock made in 1M NaOH then diluted, pH 7.4; Sigma)^10,42^ and the metabotropic glutamate receptor antagonist DL-2-Amino-3-phosphonopropionic acid (AP-3) ([1mg/kg] in PBS, stock made in 1M NaOH diluted, pH 7.4; Sigma)^10,43^; and non-selective P2-purinoreceptor antagonist pyridoxalphosphate-6-azophenyl-2’,4’-disulfonic acid (PPADs) ([25mg/kg] in PBS; Sigma).^27,44–46^ Following recording of a pre-inhibitor response, one inhibitor was delivered over one minute (devazepide and PPADS were delivered 10μL/g; KA/AP3 cocktail was delivered 20μL/g). For devazepide, infusion of the selected sugar ligand was repeated for post-inhibitor recording after a 5-8 minute incubation period. For KA/AP3 and PPADs, infusion of the selected sugar ligand was repeated for a post-inhibitor recording after a 3-5 minute incubation period.

### Data Analysis

Stimulation response was quantified as the maximum firing rate after stimulation (stimulant conditions) or during recording (baseline). Time to peak was also quantified as the time from the start of infusion to the maximum firing rate for stimulant conditions which evoked vagal firing. Each trial served as its own control by normalizing the firing rate to the pre-stimulus baseline firing rate (first 2 minutes of recording). Throughout experiments, sucrose response was used as a positive control. For all nutrient and laser stimulation conditions, data were excluded if a sucrose response was not seen throughout the recording session. Maximum firing rate, time to peak, and area under the curve were analyzed across stimulation condition.

## Immunohistochemistry

CckGFP (n = 3) mice were transcardially perfused with PBS for 3 minutes followed by 4% PFA for 3 minutes at a rate of 600µl/min. Each small intestine was harvested, opened lengthwise, rolled with the proximal end in the center, and post-fixed in 4% PFA for 3 hours. Tissue was then dehydrated in 10% sucrose for 1 hour and 30% sucrose for at least 12 hours. Samples were embedded in OCT (VWR) and stored at -80°C. Tissue was sectioned onto slides at 14μm using a cryostat. Tissue slides were post-fixed in 10% normal buffered formalin (VWR) for 10 minutes then washed in tris-buffered saline with 0.05% Tween-20 (TBST) (Sigma). SGLT1 staining was achieved by performing heat mediated antigen retrieval. Trisodium citrate dihydrate buffer (10mM in PBS, 0.05% Tween, pH 6.0; Sigma) was heated in a slide holder in a water bath to >90°C. Tissue slides were immersed for 20 minutes and then immediately placed into cool tap water then washed in TBST for 5 minutes. Tissue was blocked in 10% donkey serum (Jackson ImmunoResearch) for one hour. Tissue was then incubated with primary antibody dissolved in antibody dilution solution (PBS with 1% BSA and 0.0025% Triton-X 100) for 24 hours at 4°C then one hour at room temperature. Primary antibodies and dilutions were as follows: Rb-Anti-SGLT1 (1:100; Abcam: ab14686) and Chk-Anti-GFP (1:500; Abcam: ab13970). Following primary antibody incubation, tissue was washed in TBST then incubated with secondary antibody in antibody dilution solution for 1 hour at room temperature (Jackson ImmunoReseach: Dk-Anti-Rb-488 (1:250); Dk-Anti-Rb-Cy3 (1:250); Dk-Anti-Ck-488 (1:250)). Tissue was then washed with TBST, stained with DAPI (1:4000) for 3 minutes, washed in TBST, and mounted using Fluoro-Gel with Tris Buffer (Electron Microscopy Sciences). Imaging was done on a Zeiss 880 Airyscan inverted confocal microscope. Images were adjusted for brightness/contrast using ImageJ (Fiji). Counts are presented as the mean percentage of co-localization ± SEM.

## Dissociation and isolation of single intestinal epithelial cells

Small intestines of mice were dissociated for single cell quantitative PCR (CckGFP; n=3), calcium imaging (CckCRE_tdTomato; n=8), or *in vitro* electrophysiology (CckCRE_tdTomato; n=9) as previously described.^10^ Briefly, the proximal half of the small intestine was removed, flushed with cold PBS, and cut into sections. Tissue was shaken in 3mM EDTA in PBS for 15 minutes at 4 °C followed by a 15-minute incubation at 37°C. The epithelial layer was then mechanically detached from the muscle layer by shaking in cold PBS. Following centrifugation at 800rpm (Eppendorf 5702 RH; rotor A-4-38), the pellet was resuspended and incubated in HBSS (Gibco) with dispase and collagenase for 10 minutes at 37°C. Sample were then centrifuged (500rpm), filtered twice through a 70-μm and 40-μm filter, and resuspended in L15 media (5% FBS, 10μL/mL 10mM HEPES, 2000U/mL Pen-Strep, and 100 μL of 1700U/mL DNAse) to produce single cell suspension for further analysis.

## Single cell quantitative PCR (qPCR)

RNA isolation from single cells was performed using the Cells Direct One-Step qRT-PCR Kit (CDK kit, ThermoFisher) per manufacturer’s protocol. 5μL Lysis Buffer Mix was pipetted into each well of a 96-well plate and spun down at 500g to spread buffer. Following dissociation protocol, single cells were sorted into a U-bottom 96 well plate (Sigma) based on GFP signal using a MoFlo XDP sorter. For each mouse, 60 GFP positive cells, 30 GFP negative cells, and control wells were sorted. Control wells of 0, 10, and 100 cells were run in duplicate. Following sort, the contents of each well were pipetted into a 96-tube 0.2mL PCR plate which was then incubated in a thermocycler at 75°C for 10 minutes. After spinning to pellet, DNAase I and 10x DNAase I reaction buffer from the CDK kit were added to each well and incubated at 25°C for 5 minutes. 2μL of 25mM EDTA was added to each well, vortexed, and pelleted. The plate was then incubated at 75°C for 10 minutes to inactivate the DNAase I. Next, cDNA was synthesized and pre-amplified. Specific Target Amplification (STA) mix was made by mixing 1μL of each TaqMan probe. STA mix, superscript RT, and reaction buffer from the CDK kit were added to each sample and incubated on a thermocycler for 15 minutes at 50°C, 2 minutes at 95°C, then 20 cycles of 15 seconds at 95°C and 4 minutes at 60°C. Gene expression was then probed using the 96.96 Dynamic Array integrated fluidic circuit on a Biomark using manufacturer protocol (Fluidigm).

### Quality control

Quality of the Ct values from the Biomark output was assessed using the Fluidigm Real-Time PCR Analysis software (Fluidigm). All trials (N = 3 mice, n = 60 positive cells and 30 negative cells per mouse) were loaded simultaneously for analysis. The quality was analyzed in linear derivative mode and the quality threshold was set at 0.65 based on manufacturer’s recommendations. All curves not meeting the quality threshold were analyzed visually for smoothness (more smooth representing high quality) and entered into analysis based on comparison with passing curves. All cells not meeting quality measures or having no detected transcripts for either housekeeping gene (*Gapdh* or *Actb1*) were excluded from analysis (48 positive cells, 24 negative cells were excluded).

### Analysis

Further processing of Ct values was performed based on Stahlberg (2012).^47^ Relative quantities of cDNA molecules (RQ) were calculated using the formula RQ = 2^(Cq_cutoff_ -Cq) using a Cq_cutoff_ value of 34. The RQ value for any sample expressing no detectable transcripts for a gene was set at 0.5. All data was expressed in a log2-scale. Heat maps were generated for gene expression normalized within each gene using Qlucore Omics Explorer (Qlucore). Differential gene expression analysis (between 1) CckGFP+ and CckGFP- and 2) CckGFP+ *Slc5a1*+ and CckGFP+ *Slc5a1*-) was carried out using two-group t-test comparisons with a q-value cut-off of 0.01 (Qlucore).

## Calcium imaging

For neurons, C57BL/6J (n=3) nodose neurons were dissociated and plated as previously described.^48^ Briefly, nodose ganglia were dissected and immediately placed into 500 μL of ganglia dissociation solution containing 10 mM HEPES, 1x Glutamine, 1x N2 supplement, 1x B27 supplement, 0.5 μg/mL NGF, and 55 μg/mL of Liberase (Roche, 5401054001) in Advanced DMEM/f12. Following digestion, ganglia were rinsed twice with PBS, mechanically dissociated in dissociation solution, and filtered through a 70 μm cell strainer. They were then plated on 12mm coverslips and placed in a 37°C incubator overnight. Cells were imaged 1-2 days after plating. For enteroendocrine cells, CckCRE_tdTomato (n=8) cells were dissociated as described in the dissociation section and fluorescence sorted (BD FACSAria) selecting for tdTomato+ fluorescent cells. Cells were then plated on coverslips coated with 2.5% Matrigel (Corning #356231). Enteroendocrine cells were imaged 2-6 hours after plating. To load cells with calcium dye, cells were washed once with calcium free PBS then incubated for 45 minutes at 37°C with 5μM Fluo-4, AM and 5µM Fura Red, AM calcium dyes (Life Technologies, Carlsbad, CA) and 0.1%Pluronic F-127 (Life Technologies) in imaging buffer (120mM NaCl, 3mM KCl, 2mM CaCl_2_, 2mM MgCl_2_, 10mM HEPES, 10mM glucose). The loading buffer was then removed, and cells were washed twice with imaging buffer and placed in the dark for 15 minutes until they reached room temperature. Coverslips were then placed in the recording chamber of a Zeiss Examiner Z1 and imaged with a Hamamatsu camera (Orca-flash4.0; C11440) using the Zeiss ZEN Blue software package. Fluo-4 and Fura red emission images were obtained using 480-nm and 570-nm excitation, respectively. Images were collected at 1.5 second intervals with a 100 ms exposure time. Each recording was 210 seconds (3.5 minutes) long. Imaging buffer was continuously perfused (∼2ml/min) over the coverslips throughout the imaging session. Four stimuli were applied during each recording: 20 mM d-glucose, 2mM sucralose, 1% maltodextrin, and 50 mM KCl (at the end as an activity control). Stimuli were each delivered for 15 seconds at 30, 75, 120, and 165 seconds following the start of the recording. Following each 15 second stimulus delivery, imaging buffer was perfused to wash away the stimulus. The order of the three experimental stimuli was counterbalanced to offset any potential order effects. Each recording session concluded with the 50 mM KCl activity control.

### Analysis

Fluorescence values for each individual cell were calculated as the mean fluorescence intensity in a user-defined region of interest on Fiji software. Intracellular calcium changes were then calculated as ΔF/R (ΔF = change in fluorescence of the ratio, R; R = fluo-4/fura red) based on baseline fluorescence. Ratiometric values were then normalized to the peak KCl response. A positive response was defined as an increase in ratio >10% above baseline. Student’s paired t-test was used for a single comparison between stimuli and KCl. Significance was set at p<0.05. Statistical analyses were carried out using MATLAB software (Mathworks) and graphs were made with MS Excel.

## Neuropod cell and vagal nodose neuron co-culture

The small intestines of CckCRE_tdTomato mice were dissociated to single cell as described in the dissociation section. Cells were sorted using fluorescence activated cell sorting (BD FACSAria) selecting for tdTomato+ fluorescent cells. Cells were sorted into co-culture media (1x Glutamax, 10 10mM HEPES, 200U/ml Penicillin-Streptomycin, 1x N2 supplement, 1x B27 supplement, 10ng/ml NGF, 0.25ng/ml EGF, 50ng/ml Noggin, and 100ng/ml R-Spondin in Advanced DMEM/f12). Sorted cells were plated on 2.5% Matrigel (Corning #356231) coated 12mm coverslips at a concentration of ∼5-10k enteroendocrine cells per coverslip. Nodose neurons were dissociated from C57 BL/6J wild-type mice as described in the calcium imaging section. Neurons in media were plated evenly on up to 8 coverslips with enteroendocrine cells. Patch-clamp electrophysiology was performed 2-3 days after plating.

## Patch-clamp electrophysiology

Enteroendocrine cells and nodose neurons were co-cultured as described in the co-culture section. Co-culture coverslips were placed in the recording chamber filled with extracellular solution containing (in mM): 140 NaCl, 5 KCl, 2 CaCl2, 2 MgCl2, 10 HEPES (pH 7.4, 300-305 mOsm). CckCRE_tdTomato cells were identified by red fluorescence and neurons by their morphology and lack of fluorescence. Recordings were made using borosilicate glass pipettes pulled to ∼3.5 MΩ resistance. For voltage-clamp recordings, intracellular solution contained (in mM): 140 CsF, 10 NaCl, 0.1 CaCl2, 2 MgCl2, 1.1 EGTA, 10 HEPES, 10 sucrose (pH 7.25, 290 - 295 mOsmol). Neurons were held at -50 mV for 2 min after patching in voltage-clamp mode to stabilize cells. Membrane time constant, cell capacitance, and voltage threshold were determined using 200 ms steps from -50 mV to +20 mV in 10 mV increments. Stimuli were delivered using the SmartSquirt Microperfusion system (Automate Scientific, Inc.). Then, the SmartSquirt nozzle was brought to within 100µm of the paired enteroendocrine cell. While extracellular solution was perfusing through the chamber (∼2ml/min), either 20mM glucose or 2mM sucralose was puffed onto the cell and its axonal arbor via the SmartSquirt needle. Baseline neuronal activity was recorded in voltage-clamp for 2 min before exposure to either stimulus in alternating order for 30-60 sec, followed by a wash with extracellular solution. After exposure to stimuli, neurons were re-tested with voltage steps as described above to confirm the health of the cell.

### Data Acquisition

Recordings were carried out at room temperature using a MultiClamp 700B amplifier (Axon Instruments), digitized using a Digidata 1550A (Axon Instruments) interface, andvisualized in pClamp software (Axon Instruments). Data were filtered at 1 kHz and sampled at 10 kHz.

### Data Analysis

Cell capacitance was calculated as Cm = (τ * I_0_)/ΔE, where τ=time constant of the decaying current transient, ΔE=voltage step, and I_0_ = current transient relative to pre-pulse potential (Platzer, 2016 #123). To account for cell variability and health, max current was normalized to the cell capacitance. Data are presented as the mean ± S.E.M. in log scale. Significance was determined using a two-tailed Student’s t test with α = 0.05.

## Single cell RNA sequencing

Left (n=6) and right (n=5) nodose ganglia adult C57 BL/6J wild-type euthanized mice were dissected as described in the calcium imaging section and separated into two distinct tubes. Dissections were completed in tandem by three lab members, and all nodose were dissected within 30 minutes at which point 55 μg of Liberase (Roche) was added to each tube. Ganglia were dissociated into single cells as described in the co-culture section. The dissociated solution was then carefully laid upon a density gradient of 500 μL 12% and 500 μL 28% Percoll (Sigma) and centrifuged for 10 minutes at 2,900 x g at room temperature. Once centrifugation was complete, the top 700 μL was removed, and 700 μL of fresh dissociation solution was added. Cells were then centrifuged for 15 mins at 2,900 x g, and the final pellet was resuspended in 500 μL of PBS + 0.04% BSA and passed to the Duke University Human Vaccine Institute Sequencing Core for further processing. Capturing of single cells was performed using Chromium Single-cell 3’ v2. cDNA synthesis with PCR and library preparation were done according to the manufacturer’s guidelines. Libraries were sequenced on an Illumina NextSeq 500. The Cell Ranger pipeline version 2.1.1 was used with the mm10 mouse reference genome version 2.1.0 to convert base calls to fastq, align, map, and count genes.

### Analysis of nodose single cell sequencing data

R-package Seurat version 3.1.0 was used for the analysis.^49^ We integrated our dataset with the published atlas for the nodose ganglia as a reference.^50^ A total of 5,847 cells were sequenced with a mean of 137,352 reads and 2,817 genes detected per cell. Cells were filtered for gene content (fewer than 1000 or greater than 6000 genes detected were removed) and mitochondrial content (greater than 10%), leading to the removal of 340 cells prior to merging the left and right nodose data sets.^49^ Normalization, feature selection, scaling, and linear dimensional reduction were then performed using default parameters in Seurat. Transfer “anchors” between our data set and the reference published set were determined using the algorithm implemented in Seurat, and we classified our cells into the 24 clusters previously identified. Cluster identities were confirmed with expression levels of cluster markers, and gene expression levels across clusters was visualized using Uniform Manifold Approximation and Projection (UMAP). For glutamate and ATP receptor genes, we determined a composite gene expression of ionotropic (Gria, Grik, Grin; P2rx) and metabotropic (Grm; P2ry) family receptor genes by summing counts for the respective genes in each cell.

## Flexible gut “Illumenator” implant fabrication

### Waveguide fabrication

The step-index core/cladding flexible polymer waveguide was fabricated using the Thermal Drawing Process (TDP) starting from a macroscopic polymer preform (template).^32,33,51^ The preform was assembled by inserting a polycarbonate (PC) rod (1/8” diameter, McMaster-Carr) into a poly-methyl methacrylate (PMMA) tube (1/4” outer diameter and 1/8” inner diameter; US Plastics Corp.) and then consolidating the rod-in-tube assembly at 170°C in a vacuum oven. The resulting preform was drawn into meters long fiber in a custom-built fiber drawing tower at a temperature of 270°C. The lateral dimensions of the preform were reduced by 30 times to produce a microscopic (220-230μm diameter) PC/PMMA core/cladding optical waveguide.

### Physical characterization of flexible polymer Waveguide

Optical transmission loss of the fibers was quantified by coupling the fibers to a diode-pumped solid state (DPSS) laser (Laserglow, 50-mW maximum output, wavelength λ = 473 nm) via ferrules and the light output was measured by a photodetector (S121C, 400–1,100 nm, 500 mW, Thorlabs) attached to power meter (PM100D, Thorlabs). Optical transmission was quantified for a range of fiber lengths (1–10 cm), bending angles (0°, 90°, 180° and 270°), and radii of curvature (0.5, 1, 2.5, 5, 7.5, 10, 12.5 and 15 mm).

### Gut implant fabrication

To optically couple as-drawn fibers to a light source, 9-10cm long fibers were inserted into a 10.5mm-long, 2.5-mm-diameter, 231μm bore size ceramic ferrule (Thorlabs) and affixed with optical epoxy (Thorlabs). The ferrule edge was then polished using a Thorlabs fiber polishing kit. Fiber was then threaded through ∼7.5cm of micro-renathane tubing (BrainTree Scientific) to provide structural stability for tunneling. The proximal ∼6.75cm of the tubing closest to the ferrule was opacified with liquid electrical tape (Starbrite) to reduce non-specific activation of Cck expressing cells in the skin (Fig. 6a). The final length of the device was ∼9.25cm including the ferrule; ∼1.5 cm of the device could be illuminated and ∼0.75cm of fiber extended beyond the tubing. The average power recorded from the device tip was measured using a photodetector (S140C, 250–1100 nm, 500 mW; Thorlabs) attached to a power meter (PM100D, Thorlabs). Average power output at the end of the PC/PMMA fiber with a 5V, 40Hz, 532nm laser input was 13.3 ± 0.107 mW/cm^2^ before implantation and 10.2 ± 1.43 mW/cm^2^ 4-8 weeks after implantation following completion of behavioral studies (n = 11).

## Gut fiberoptic implantation surgery

Adult CckCRE_Halo-YFP mice and adult CckCRE_ChR2-tdTom were single-housed and acclimated to behavioral cages (TSE PhenoMaster) one week prior to surgery. Mice were anesthetized with isoflurane (1-3% in oxygen). A 2 cm incision was made from the xiphoid process diagonally to left-mid clavicular line. The peritoneal cavity was accessed, and the stomach extra-corporealized for implantation. A purse string suture was made in the gastric antrum, avoiding blood vessels. A small incision was made in the stomach within the suture, and a gavage needle was used to dilate the pylorus. The distal end of the device was threaded into the proximal duodenum so that the illuminated region of the device was in the proximal small intestine (Fig. 4e). The purse string stitch was tied to secure the device in the intestine. The opacified portion of the device was tunneled to the base of the skull. The peritoneum and overlying skin were sutured. The device exited the tunnel at the base of the skull and was skull mounted; skull mounting was required to maximize longevity of the implant. For maximal adhesion, the skull was etched with a razor blade and a thin layer of Metabond cement (Clear L-powder S399 + catalyst; Metabond) was applied. Then, the Metabond layer was etched and the device attached using standard dental cement (Stoelting #51458). Mice recovered for at least 5 days or until normal feeding behavior and activity returned.

## Duodenal catheter surgery

Adult C57BL/6J wild-type mice were surgically implanted with catheters into the duodenum with a similar procedure as previously described.^4,52^ Mice were anesthetized with isoflurane (1-3% in oxygen). A 2 cm incision was made from the xiphoid process diagonally to the left-mid clavicular line. The peritoneal cavity was accessed, and the stomach extra-corporealized for catheter implantation. Micro-renathane tubing (BrainTree Scientific) with one silicone ball (Home Depot) at implantation end was inserted into a small incision made in the stomach within a purse string suture. The distal end of the catheter was threaded into the proximal duodenum, and the silicone ball was sutured inside the stomach to keep the catheter in place. The other end of the catheter was tunneled to the back and directed out in the small intrascapular incision. This end was secured in place with surgical mesh. The proximal end of the catheter was sealed with a metal cap. Mice were then singly housed and recovered for at least 5 days until normal feeding behavior and activity returned.

## Phenotyping equipment

All optogenetic behavior experiments were performed in a PI managed husbandry system. Animals were housed in a custom-built PhenoMaster behavioral phenotyping system (TSE Systems Inc. Chesterfield, MO). The PhenoMaster was programmed (software version 6.6.9) to automatically maintain a light cycle (0700 lights on; 1900 lights off), temperature control (22°C), and humidity control (40%). The PhenoMaster holds 12 clear cages in which animals were singly housed. Cages were industrially washed and bedding (ALPHA-dri) replaced weekly. Animals were provided with standard mouse chow (Purina 5001) and reverse osmosis water *ad libitum* unless fasted for a choice assay. All cages also housed an enrichment device, which also served to weigh the animals. Food hopper, water bottle, and weigh container were attached to weight sensors (TSE). Food intake, water intake, and weight were automatically measured every 5 seconds to the nearest 0.01g. For drinking measurements, a 10 second smoothing interval with a maximum raw analog-to-digital conversion counts difference of 40,000 was permitted. For weight measurements, a 15-second smoothing interval with a 15g threshold and a maximum raw analog-to-digital conversion counts difference of 1,000,000 was permitted. Intake was measured every 5 seconds and binned every minute for analyses unless otherwise indicated. Animal activity was determined by beams crossed in the x and y planes and was collected with a 100 Hz scan rate. For optogenetic stimulation experiments, custom PhenoMaster software drove scheduled TTL pulses which triggered laser on/off. For optogenetic experiments, TTL signals were set to be triggered every 3 minutes. Each cycle included one-minute on with 40 Hz, 5V pulse at 20% duty cycle, and two-minutes off. Each experimental session with laser stimulation began with 1-minute laser on. Following experiments, raw data was downloaded from the Phenomaster software and analyzed using MATLAB software (Mathworks). Unless otherwise indicated, all activity, food intake, and water intake measurements were binned in 1-minute intervals for analysis. Data was corrected for minor fluctuations by only permitting a monotonically increasing function for both food and water intake: values that represented positive food intake were replaced by the most recent value.

### Plexiglass cage manufacturing

The choice assay paired with intraluminal drug delivery occurred in in-house manufactured clear plexigass cages. Clear plexigass (Home Depot Model # acr0802448; 0.08 x 24 x 48 inch) was manufactured into cages with an 8 x 8 inch base and four 10 inch tall walls, secured with clear silicone (Loctite waterproof sealant). The walls were snap-cut by hand in our laboratory or in the Duke Engineering machine shop. The top of the cage was open. To allow mice to move around the cage freely, a custom swivel arm (TSE), which introduced the tubing to attach to each catheter, was secured on one wall with a custom 3-D printed device. This device was essentially a tube which snuggly fit around the metal swivel arm to hold it upright and was super glued to one side of the cage.

## Choice assay

Mice were given free access to 300mM sucrose and 15mM sucralose for 24 hours in the home cage to control for neophobia. During 24-hour access, mice had *ad libitum* access to food and water, though water intake was negligible; implanted mice were not connected to patch cables. For each subsequent choice assay, at the start of the dark cycle (1900), mice were placed in a cage with fresh bedding and restricted of food and water either in the Phenomaster or plexiglass cages. One hour after onset of dark cycle (2000), 300mM sucrose and 15mM sucralose became available for free consumption for 1 hour. Concentrations were selected based on prior studies showing iso-sweetness.^53,54^ The side of the sucrose and sucralose solutions was swapped each test to control for side preferences. In order to advance to optogenetic or pharmacologic inhibition, mice were required to display a stable preference for sucrose, defined as > 66% sucrose preference in two consecutive tests, not varying by more than 15% across both tests. For five mice who did not display a clear preference by the seventh test, or displayed a clear side preference by the third test, were re-exposed for 24 hours as above. Average number of test days to advance to inhibition was 5.24. Following each test session, mice were disconnected when appropriate (optogenetic and intraduodenal infusion tests) and given *ad libitum* access to food and water. The start of all test sessions was separated by at least 48 hours.

### Optogenetic inhibition

Implanted CckCRE_Halo-YFP mice (final n = 8, n = 5 male/3 female) and their wild-type littermates (final n = 5, n = 3 male/2 female) underwent baseline choice assays as described in choice assay section. Testing occurred in our TSE PhenoMaster apparatus. During all baseline and experimental assays, mice were connected to the laser using a custom swivel arm (TSE) coupled to a rotary joint patch cable (ThorLabs, #RJPFF2) for free movement at the time of dark onset for consistency and acclimation. Once a stable preference was established, each mouse underwent two experimental conditions: 532nm (silencing) laser and 473nm (control) light, followed by a repeated baseline. Order of laser wavelength was counterbalanced to control for order effect. During photo-stimulation conditions, the Phenomaster delivered a TTL pulse for laser stimulation: 1 minute on (5V, 40Hz, 20% duty cycle) and 2 minutes off for the full hour. The following analyses were assessed: minute-to-minute sucrose and sucralose consumption, preference for sucrose relative to the total amount of volume consumed, motor activity, and food/chow intake over 24 hours following assay. Fiberoptic placement and power output was confirmed at the end of the study. Only mice who completed all four tests and whose device had appropriate power/placement were included in analysis. Four mice were excluded for the following reasons: 1 mouse for low power, 3 mice due to a strong side preference.

### Pharmacologic inhibition with intraperitoneal injections

C57BL/6J wild-type mice (for CCK-A inhibition, final n = 4, n = 3 male/1 female); for IP glutamate receptor inhibition, final n = 4, n = 2 male/2 female) underwent baseline choice assays as described in choice assay section. During acclimation period, wild type mice received intraperitoneal injections of PBS to acclimate them to the procedure. Once a stable sucrose preference was established, each mouse underwent two experimental conditions: drug and vehicle. Order of injection was counterbalanced. Separate cohorts of wild type mice were used to test the effect of the CCK inhibitor devazepide (Bachem) and the glutamate receptor inhibitor cocktail KA/AP3 (Sigma).^42,43^ Testing occurred in our TSE PhenoMaster apparatus as described in the optogenetics section or plexiglass cages. For experiments in plexiglass cages, solution intake was measured to the nearest 0.1mL manually every 5 minutes from 5 mL serological pipettes fashioned as sippers. For CCK-A inhibition, devazepide was delivered 2mg/kg (10μL/g) dissolved in 5% DMSO in PBS. Devazepide or vehicle (5% DMSO in PBS) was injected intraperitoneally 30 minutes prior to choice assay based on prior reports.^17^ For glutamate receptor inhibition, stock solutions of kynurenic acid and AP-3 were made in 1M NaOH and experimental concentration were diluted in PBS, pH 7.4. KA/AP-3 were delivered at 150μg/kg and 1mg/kg, respectively, and delivered 10μL/g. KA/AP3 cocktail or vehicle (equivalent volume 1M NaOH in PBS, pH 7.4) was delivered intraperitoneally immediately before testing. Two mice were excluded from further study due to a strong side preference.

### Pharmacologic inhibition with intraluminal injections

C57BL/6J wild-type mice (final n = 4, n = 3 male/1 female) were implanted as described in the duodenal catheter surgery section and underwent baseline choice assays as described in choice assay section. Catheters were flushed with 0.05 mL PBS daily and immediately before the one-hour food and water deprivation (dark onset). Testing occurred in our plexiglass cages with syringes (solution intake measured manually every 5 minutes). Immediately prior to the one-hour test session, mice were attached to micro-renathane tubing which delivered drug, vehicle, or control solutions via an infusion pump (Fusion 200, Chemyx). The infusion pump was started, which delivered 0.4 mL of the appropriate solution over the one-hour test session (flow rate was 0.0066 mL/minute). PBS solutions were delivered intra-intestinally during baseline tests. For glutamate receptor inhibition, KA/AP3 cocktail (15 ng KA and 0.1 μg AP-3) or vehicle (1M NaOH in PBS, pH 7.4) was delivered intra-intestinally in 0.4 mL. Catheter placement was confirmed at the end of the study by visual inspection of the small intestine following the infusion of a dye solution through the catheter. One mouse was sacrificed before the final baseline test out of concern for stability of the implant. However, the catheter was confirmed to be in the duodenum, and its data were included. One mouse was excluded from further study due to a strong side preference.

## One-bottle assay

CckCRE_ChR2 mice (final n = 4, n = 2 male/2 female) were acclimated to the PhenoMaster and implanted with flexible fiberoptic implants as described above (section: *Gut fiberoptic implantation surgery)*. Implanted mice were given free access to sucrose [300mM] or sucralose [15mM] separately for 24 hours in the home cage to control for neophobia; exposures were separated by at least 48 hours. During 24-hour access, mice had *ad libitum* access to food and water, though water intake was negligible; implanted mice were not connected to patch cables during exposure. For each one-bottle assay, 5 minutes before the start of the dark cycle, access to food and water was closed and mice were attached to the rotary joint patch cable (ThorLabs, #RJPFF2) in the Phenomaster; mice remained in the home cage. At the start of the dark cycle, 300mM sucrose OR 15mM sucralose became available for free consumption for 1 hour. The side of the solution was the same for all studies. Following each test session, mice were disconnected and given *ad libitum* access to food and water. The start of all test sessions was separated by at least 48 hours.

Each mouse underwent two baseline conditions (sucrose [300mM] no laser, sucralose [15mM] no laser) followed by four experimental conditions: sucrose + 473nm (activating), sucrose + 532nm (control), sucralose + 473nm, and sucralose + 532nm. Order of experimental conditions were counterbalanced to control for order effect. During photo-stimulation conditions, the Phenomaster delivered a TTL pulse for laser stimulation based on intake as follows: for every 0.01g of liquid consumed, the mice received 5 seconds of laser stimulation (5V, 40Hz, 20% duty cycle). Fiberoptic placement and power output was confirmed at the end of the study. Only mice who completed all six tests and whose device had appropriate power/placement were included in analysis. One mouse was excluded from analysis for low device power at sacrifice.

## Statistics and reproducibility

We performed statistical analyses using JMP Pro Software (Version 14, SAS), unless otherwise indicated. For comparison of means, ANOVA was used and post-hoc testing was performed when applicable. For vagal nerve recordings, we performed Tukey HSD post-hoc testing. For behavior studies, we used a repeated measure ANOVA to account for each subject and followed with post-hoc paired student’s t-tests. For other studies, comments on statistical tests performed is included throughout methods and in figure legends. All error bars and shaded regions represent standard error of the mean (SEM) unless otherwise indicated. Sample size was not pre-determined using power analyses. All behavioral studies were counterbalanced across gender and sex to control for variables including position in cage, order effect, and handedness. Experimenters were not blinded to treatment condition, genotype, or outcome.

## Extended Data

**Extended Data Video 1 | The illumenator.** A conventional stiff, silica fiber (left) punctures an agar [1.5%] membrane, but the flexible PC/PMMA fiber (right) does not.

**Extended Data Figure. 1.**
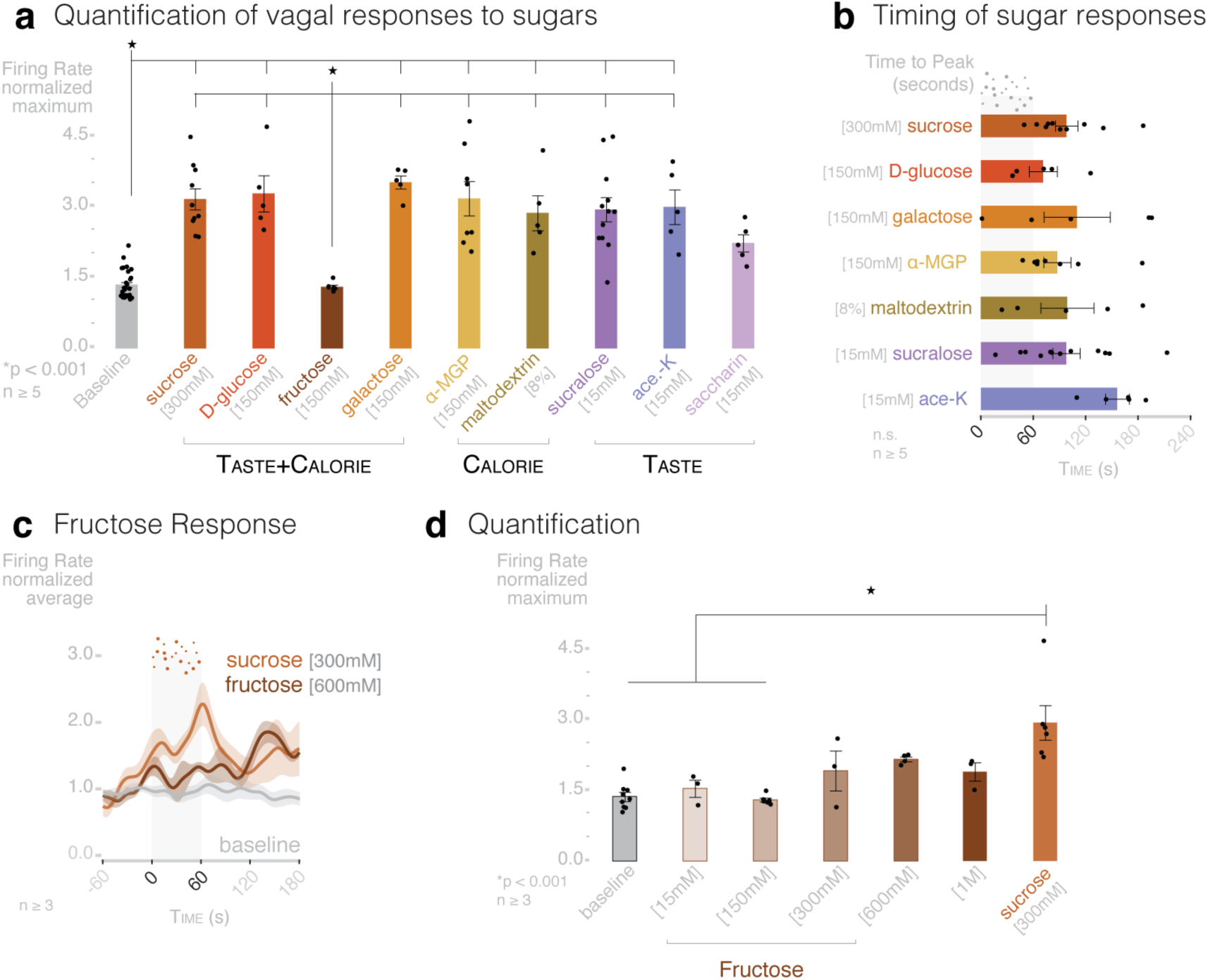
Caloric and non-caloric sugars elicit a rapid vagal response. Additional data relating to Fig 1. **a**, Whole sugars (sucrose [300mM], D-glucose [150mM], fructose [150mM], galactose [150mM]), non-sweet sugars (alpha-methylglucopyranoside (α-MGP) [150mM], and maltodextrin [8%]), and non-caloric sugars (sucralose [15mM], acesulfame K (ace-K) [15mM], and saccharin [15mM]) were intraluminally delivered while recording vagal activity. All sugars elicited significant vagal firing compared to baseline except for fructose and saccharin in wild-type mice. (n ≥ 5 mice per group; *p < 0.001, ANOVA with post hocTukey’s HSD test). **b**, Time to peak vagal firing rate was similar across all stimuli that elicited a response (n ≥ 5 mice per group; n.s. by ANOVA with post hocTukey’s HSD test). **c**, Normalized vagal traces for sucrose [300mM], fructose [600mM], and baseline in wild-type mice. **d**, High concentrations of fructose (600mM, 1M) elicit a blunted, but not significant, vagal response. (n ≥ 3 mice per group; statistics by ANOVA with post hocTukey’s HSD test). Vertical bars indicate infusion period; all shaded regions and error bars represent SEM.

**Extended Data Figure. 2.**
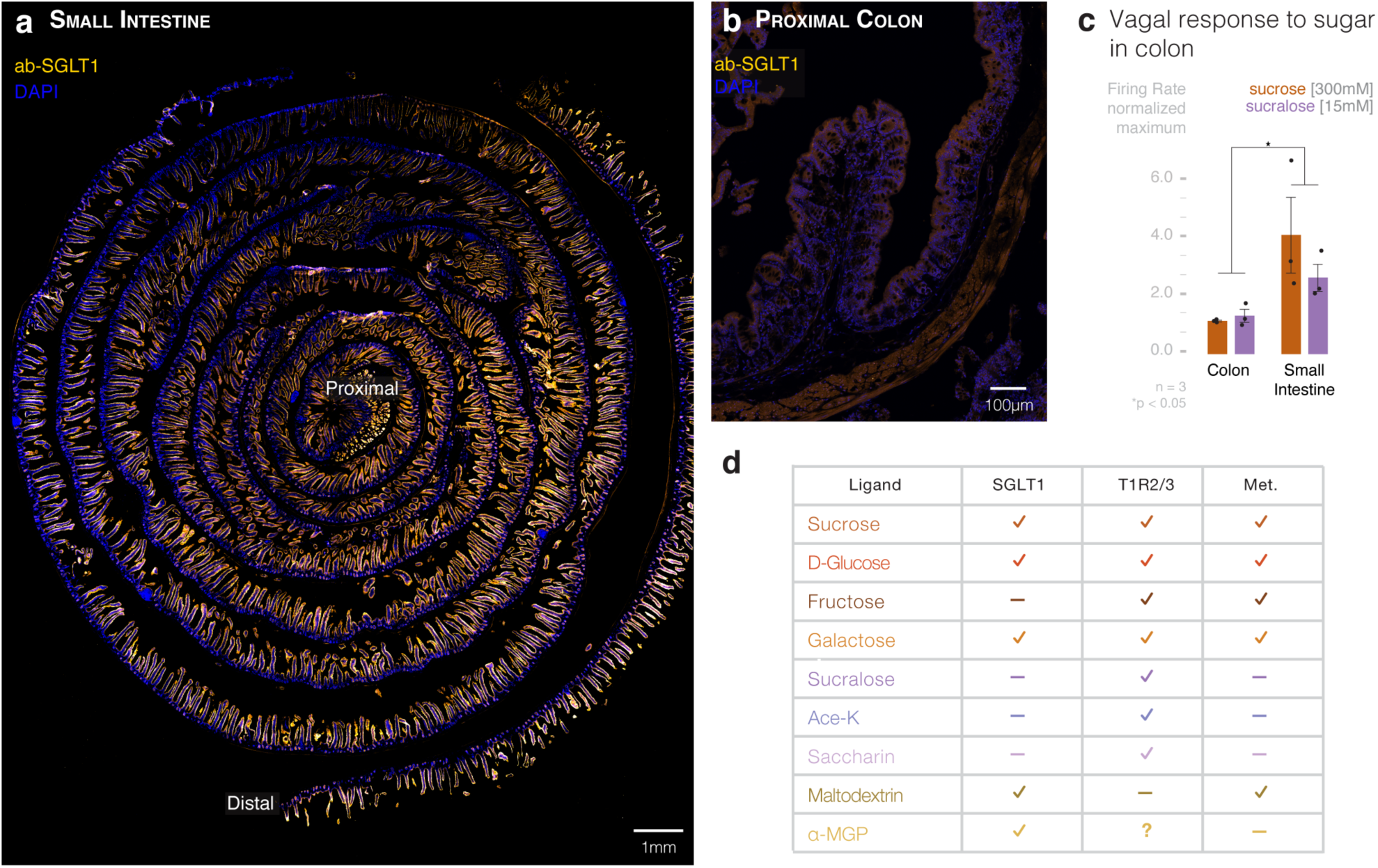
Intestinal SGLT1 is expressed consistently along the small intestine but minimally in the colon. **a**, A cross-section of the entire length of the small intestine shows consistent SGLT1 (yellow) expression across the length (proximal = center). Scale bar = 1mm. **b**, Minimal SGLT1 expression is visualized in the proximal colon. Scale bar = 100μm. **c**, Sucrose [300mM] and sucralose [15mM] infused intraluminally into the proximal colon do not elicit a vagal response (n ≥ 3 mice per group; *p < 0.05, ANOVA with post hocTukey’s HSD test; error bars indicate SEM). **d**, Caloric and non-caloric sugars act on different receptors (SGLT1 and T1R2/3) and are differentially metabolized (Met.).

**Extended Data Figure. 3.**
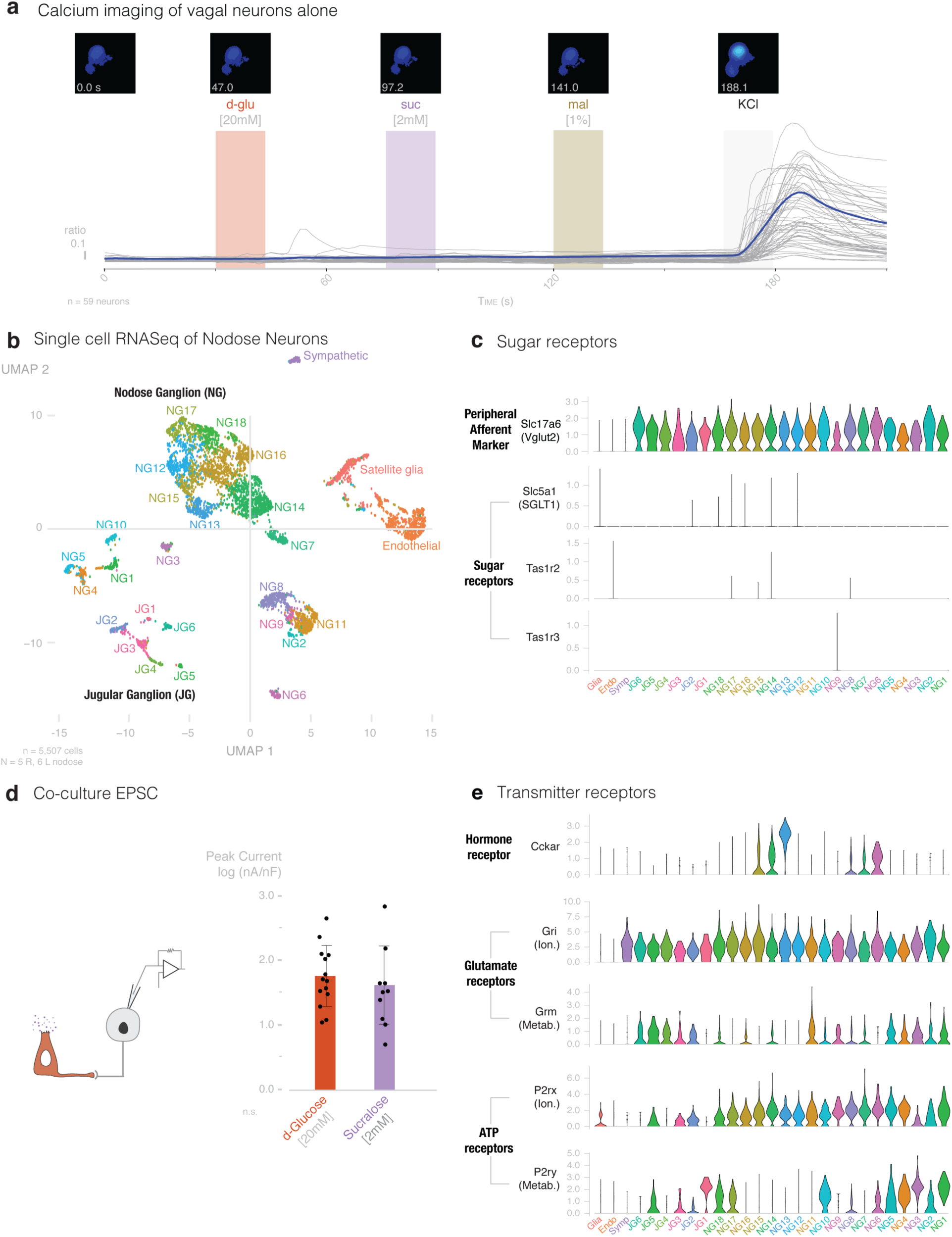
Vagal nodose neurons have transcripts for both glutamatergic and purinergic neurotransmission. **a**, Vagal nodose neurons cultured alone do not respond to D-glucose [20mM], sucralose [2mM], or maltodextrin [1%] (n=59 neurons). **b**, Single cell transcriptomic data projected onto the vagal nodose atlas showing 18 Nodose (NG) and 6 Jugular (JG) clusters**. c**, Violin plots showing transcripts for *Slc5a1* (SGLT1), *Tas1r2*, and *Tas1r3* are not expressed in vagal neurons. VGLUT2 (*Slc17a6)*, a peripheral afferent marker, is found ubiquitously in nodose and jugular ganglion neurons. **d**, Additional data related to Figure 2f. In co-culture patch-clamp electrophysiology, the peak excitatory post-synaptic currents (EPSCs) elicited in connected nodose neurons was not significantly different between the application of D-glucose [20mM] and sucralose [2mM]. (n = 18 pairs)**. e**, Several NG clusters (NG6-8, 13-5) show expression of *Cckar* transcripts. Ionotropic (Ion.) and metabotropic (Metab.) glutamatergic receptors and P2rx (ionotropic) and P2ry (metabotropic) purinergic receptors are also expressed in several NG and JG clusters. (for b, c, e: n = 5,507 cells; N = 5 right, and 6 left murine nodose ganglia).

**Extended Data Figure. 4.**
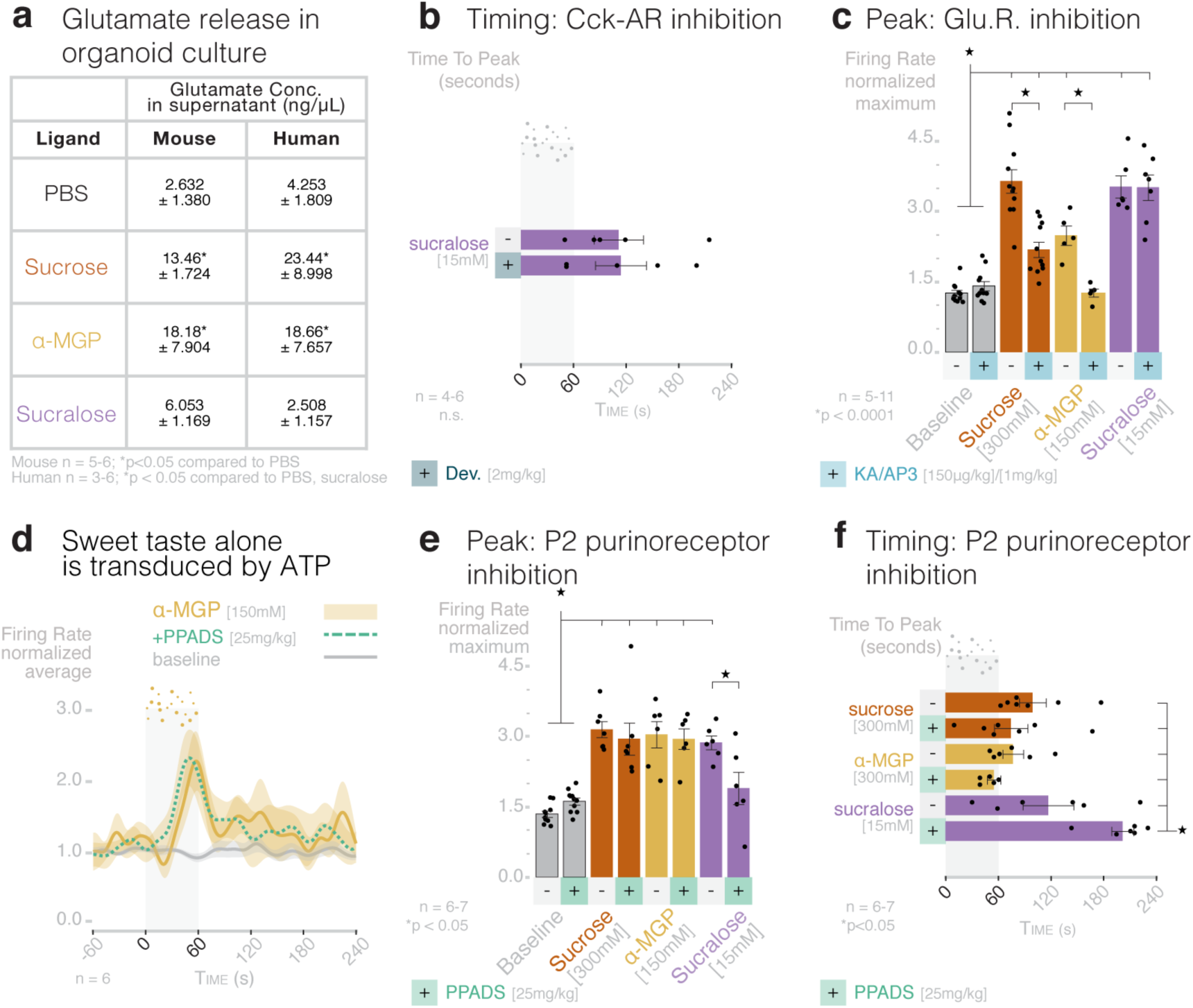
Neuropod cells use different neurotransmitters to distinguish caloric sucrose from non-caloric sweeteners. **a**, Additional data related to Figure 3b. Quantification of glutamate concentration in supernatant for mouse and human organoids stimulated with PBS, sucrose [300mM], αMGP [150mM], and sucralose [15mM] (mouse: n=5-6 plates, N=3 mice, *p<0.05 by Student’s t-test compared to PBS; human: n=3-6 plates, N=1 human sample, *p<0.05 by Student’s t-test compared to PBS, sucralose). **b**, Additional data related to Figure3c. Cck-A inhibition with devazepide (dev.; 2mg/kg) did not significantly change time to peak vagal firing rate in response to sucralose [15mM]. **c-f**, Additional data related to Figure3c-e. **c**, Quantification of peak vagal firing rate to sucrose [300mM], sucralose [15mM], and αMGP [150mM] before (-) and after (+) inhibition with the glutamate receptor inhibitors kynurenic acid (KA) [150μg/kg] and AP3 [1mg/kg] (n=5-11 mice, *p<0.0001 ANOVA with post hoc Tukey’s HSD test). **d**, Inhibition of non-selective P2 purinoceptors with PPADS [25mg/kg] did not attenuate the vagal response to αMGP [150mM]. **e**, Quantification of peak vagal firing rate to sucrose [300mM], sucralose [15mM], and αMGP [150mM] before (-) and after (+) inhibition with PPADS (n=6-7 mice, *p<0.05 ANOVA with post hoc Tukey’s HSD test). **f**, Non-selective P2 purinoceptor inhibition with PPADS prolonged time to peak vagal firing rate to sucralose [15mM] but not sucrose [300mM] or α-MGP [150mM] (n=6-7 mice, *p<0.05 ANOVA with post hoc Tukey’s HSD test). Vertical bars indicate infusion period; all shaded regions and error bars represent SEM.

**Extended Data Figure. 5.**
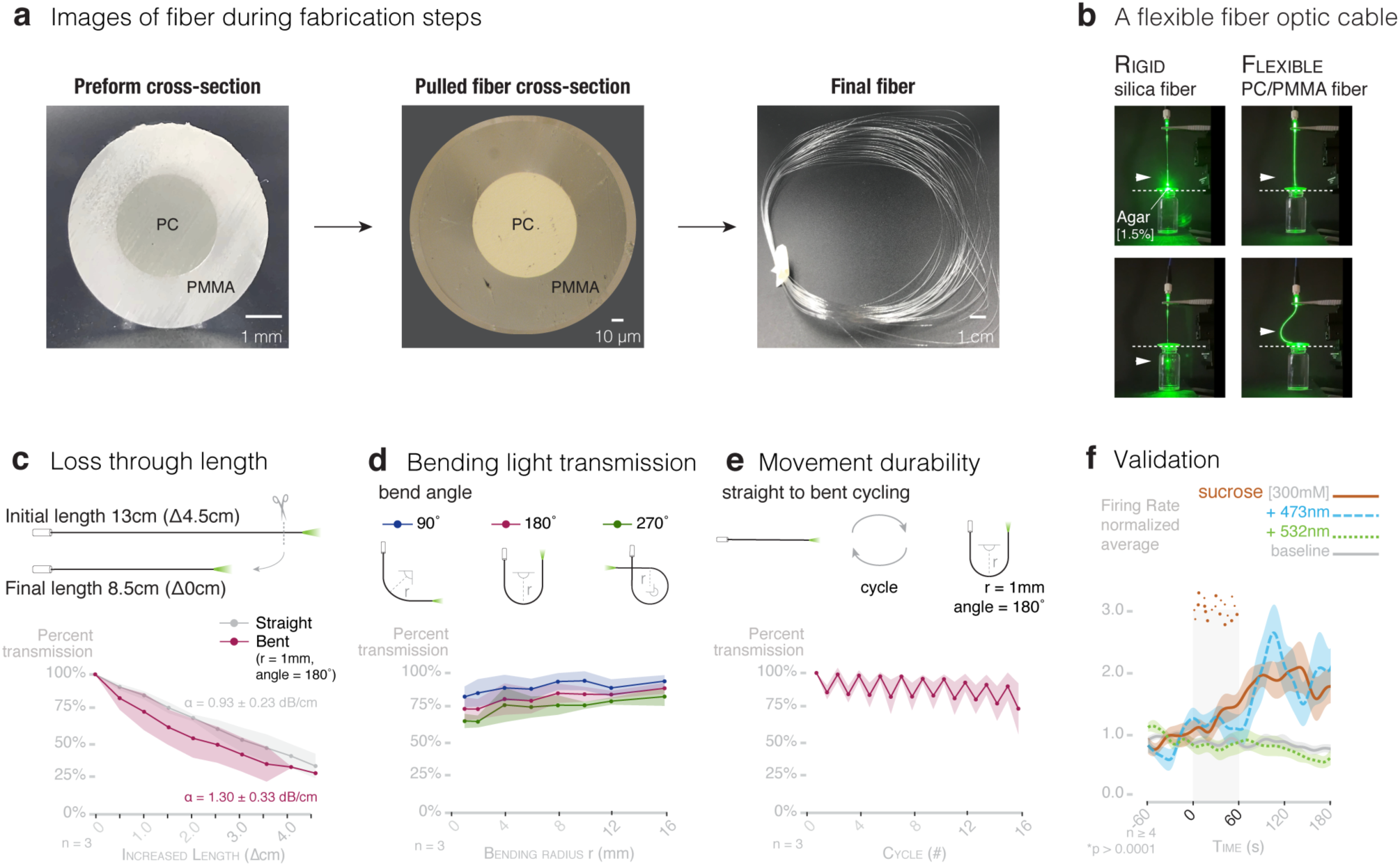
The illumenator. **a**, Cross section images of PC/PMMA core/cladding preform, as-drawn PC/PMMA flexible fiber, and photograph of ∼50m fiber bundle. **b**, A novel flexible fiberoptic for gut intraluminal optogenetics in awake and behaving mice. A conventional rigid silica fiber punctures an agar [1.5%] membrane, but the flexible fiber does not. **c**, Light transmission loss through the length of fiber. Light transmission for straight and bent (angle = 180°, radius of curvature = 1mm) flexible waveguides using the cut-back method. Percentage of light output from shortest length (Δ0cm). Loss coefficients were calculated to be 0.93 dB/cm and 1.30 dB/cm for straight and bent fibers, respectively (shaded regions represent SD). **d**, Light transmission for fibers bent at angles 90°, 180°, and 270° for varied radii of curvature. Plotted as percentage of output from a straight fiber (shaded regions represent SD)**. e**, Light transmission for flexible waveguides during cyclic bending at 180° with a 1 mm radius of curvature (odd cycles = straight, even = bent). Plotted as percentage of output from initial position (cycle = 0) (shaded regions represent SD). **f**, Four weeks after implantation of the flexible fiberoptic in CckCRE_NpH3 mice, vagal recordings show its efficiency in silencing neuropod cells with 532nm, but not 473nm, laser. Vertical bar indicates infusion period; shaded regions represent SEM (n≥4 mice; *p < 0.0001 comparing maximum firing rate, ANOVA with post-hoc Tukey’s HSD test).

**Extended Data Figure. 6.**
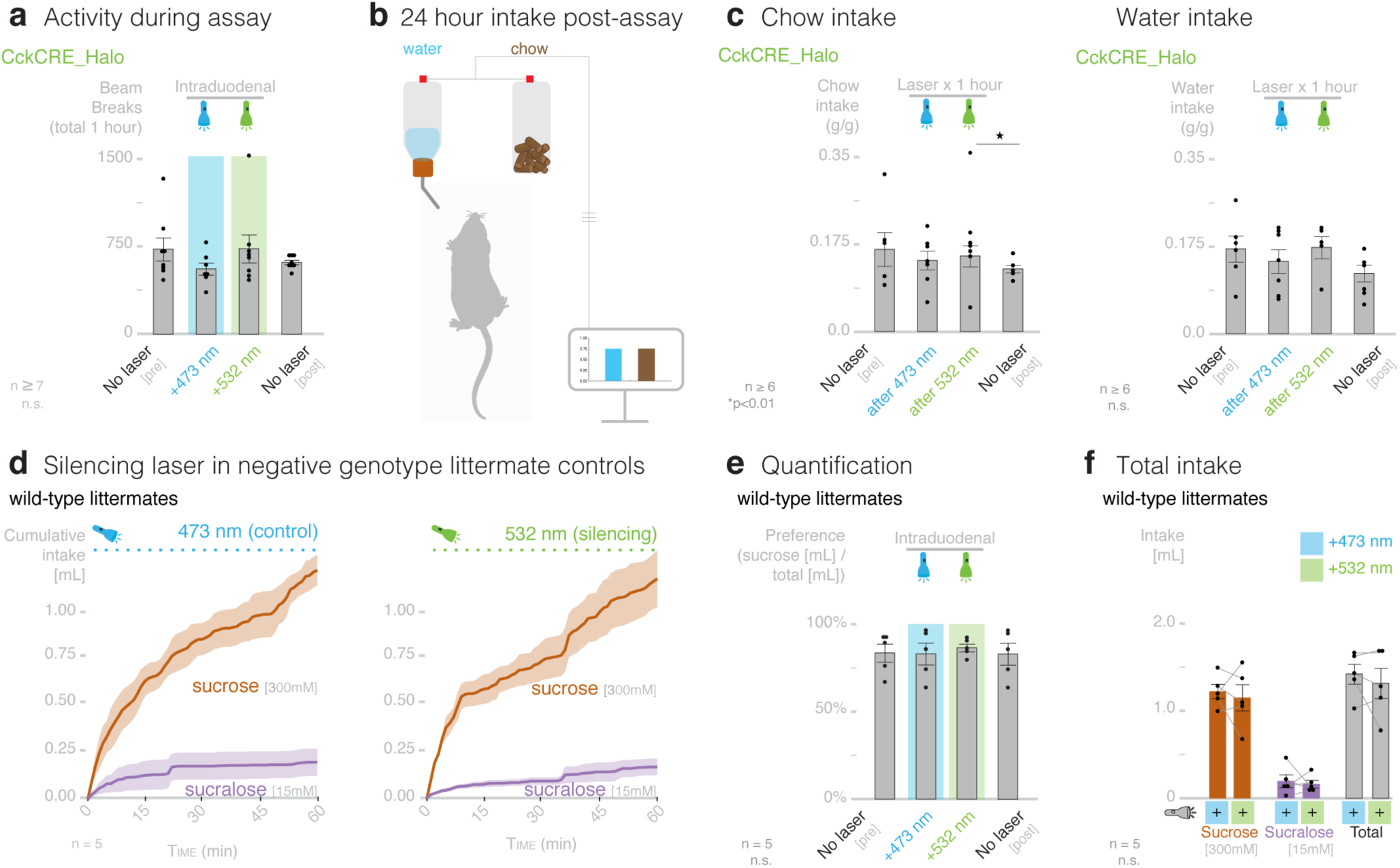
Silencing neuropod cells does not cause malaise. **a**, Activity during assay is unchanged by 473nm (control) and 532nm (silencing) in Cck_Halo mice (n≥7, n.s. (p>0.05) by repeated measures ANOVA). **b**, Water and chow consumption were measured during the 24 hours following the choice assay. During this time, mice moved freely and were not connected to laser. **c**, 532nm silencing laser applied for 1 hour during choice assay did not affect 24-hour intake of chow or water compared to 473nm control laser applied for one hour during choice assay (n≥6, *p<0.01 by repeated measures ANOVA with paired t-test post-hoc analysis). **d**, Average traces showing sucrose [300mM] and sucralose [15mM] consumption during 473nm (control, left) laser and 532nm (silencing, right) laser inhibition in genotype-negative littermate controls of CckCRE_NpH3 mice. **e**, Preference for sucrose over sucralose is unchanged by 532nm or 473nm laser in genotype-negative littermate control mice. **f**, 532nm or 473nm does not affect sucrose, sucralose, or total intake in control mice (n = 5 littermate controls; n.s. (p>0.05) by repeated measures ANOVA).

**Extended Data Figure. 7.**
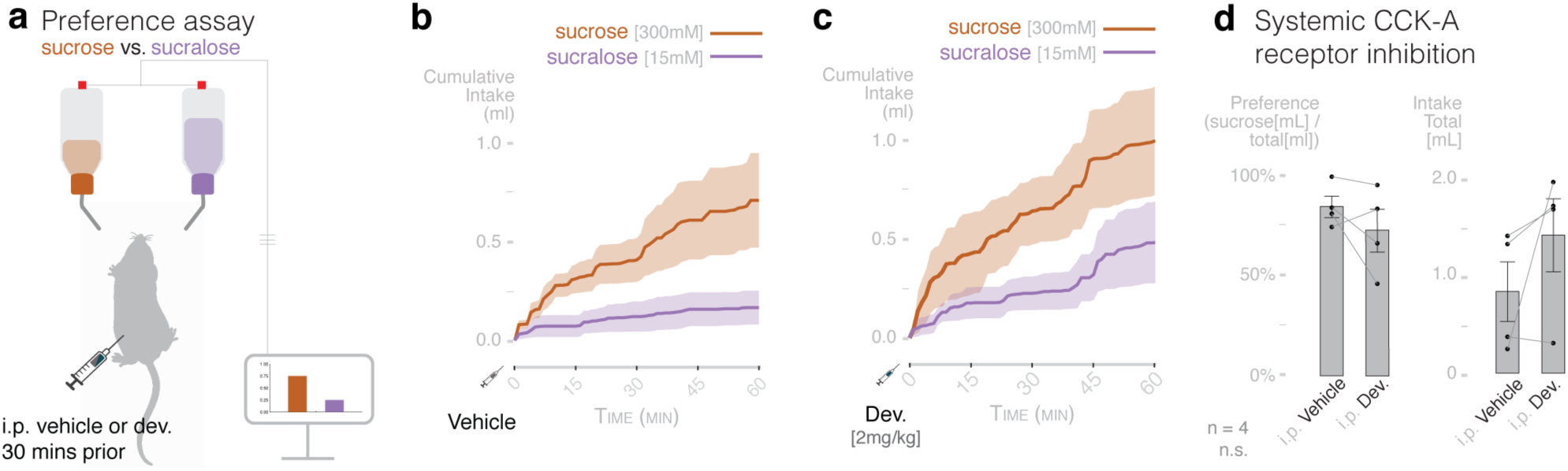
Intraperitoneal cholecystokinin receptor inhibition does not eliminate preference for sucrose. **a**, Mice are given a two-bottle preference test between sucrose [300mM] and sucralose [15mM] for 1 hour. Drug or vehicle is administered intraperitoneal 30 minutes prior to assay. **b-c**, Average traces showing sucrose [300mM] and sucralose [15mM] consumption after intraperitoneal injection 30 minutes prior to solution access of **b**, vehicle (PBS + 5% DMSO) or **c**, CCK-A receptor inhibitor devazepide ([2mg/kg] in 10μL/g mouse in 5% DMSO in PBS) in wild-type mice. **d**, Preference for sucrose over sucralose is unchanged by devazepide compared to vehicle in wild-type mice (n = 4, n.s. p = 0.6257 on repeated measures ANOVA). Total intake trends toward increasing with Cck-A receptor inhibition compared to vehicle control (n = 4, n.s. p = 0.2353 on repeated measures ANOVA).

**Extended Data Figure. 8.**
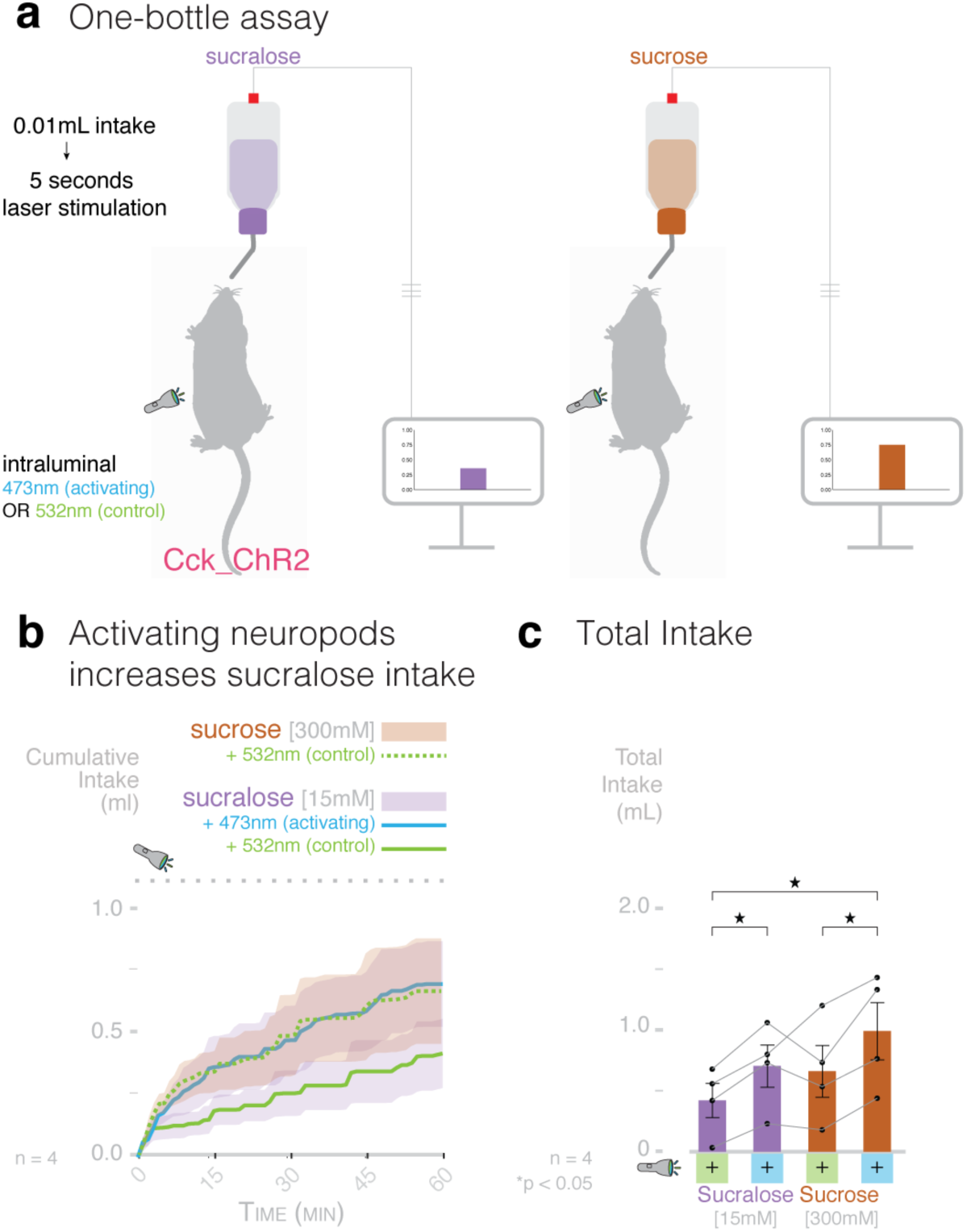
Optogenetic stimulation of neuropod cells increases consumption of caloric and non-caloric sugars in a one-bottle test. **a**, Design: CckCRE_Channelrhodopsin (Cck_ChR2) mice are given a one-bottle intake test of sucrose [300mM] or sucralose [15mM] for 1 hour with 532nm (control) or 473nm (activating) laser. Laser stimulation is paired to solution consumption: for every 0.01mL intake, mice receive 5 seconds of intraluminal stimulation at 40Hz. Solution*laser conditions were counter-balanced to control for order effect. **b**, Average traces showing consumption of sucrose [300mM] + 532nm control laser, sucralose [15mM] + 473nm activating laser, and sucralose [15mM] + 532nm control laser in Cck_Chr2 mice. **c**, Stimulation of duodenal neuropod cells with 473nm increases consumption of both sucrose [300mM] and sucralose [15mM]. Neuropod activation with 473nm increases consumption of sucralose to consumption of sucrose with control laser (n = 4; *p<0.05, repeated-measures ANOVA with post-hoc paired t-test). Shaded regions or error bars indicate SEM.

